# Integrative epigenomic and functional characterization assay based annotation of regulatory activity across diverse human cell types

**DOI:** 10.1101/2023.07.14.549056

**Authors:** Tevfik Umut Dincer, Jason Ernst

## Abstract

We introduce ChromActivity, a computational framework for predicting and annotating regulatory activity across the genome through integration of multiple epigenomic maps and various functional characterization datasets. ChromActivity generates genomewide predictions of regulatory activity associated with each functional characterization dataset across many cell types based on available epigenomic data. It then for each cell type produces (1) ChromScoreHMM genome annotations based on the combinatorial and spatial patterns within these predictions and (2) ChromScore tracks of overall predicted regulatory activity. ChromActivity provides a resource for analyzing and interpreting the human regulatory genome across diverse cell types.

## Background

Transcriptional regulation of gene expression is controlled by a large set of regulatory elements distributed across the genome [1–3]. Identifying and predicting regulatory elements is important to advancing our understanding of cellular processes and gaining insight into the genetic basis of common diseases [1, 4, 5].

Epigenomic data, such as maps of histone modifications, histone variants and chromatin accessibility, have been powerful resources for the identification of candidate regulatory elements within the genome [4, 6–9]. Such data is now available across hundreds of different cell or tissue types based on the efforts of large consortium projects [7, 9, 10] as well as contributions from individual labs [6, 11]. Maps of chromatin marks enabled the prediction of regulatory elements in hundreds of cell types, often through unsupervised approaches such as calling peaks on single marks [12] or the identification of combinatorial and spatial patterns of multiple marks using chromatin state models [13–16].

However, despite its extensive utility, unsupervised integration of chromatin marks does not take advantage of information from functional characterization assays to potentially better predict regulatory regions. Functional characterization assays complement chromatin marks by enabling direct testing of genomic regions for regulatory activity in high-throughput [17–20] by either incorporating sequences of candidate regulatory elements into cells via plasmids or by manipulating or interfering with the genome itself using lentiviral integrases or CRISPR-based technologies [3]. Plasmid-based assays [21], such as barcoded Massively Parallel Reporter Assays (“MPRAs”) [22, 23] or Self-Transcribing Active Regulatory Region Sequencing (STARR-seq) assays [24], typically measure the expression of a reporter gene on a plasmid containing the candidate regulatory element, serving as an indicator of the expression level that it is likely to induce in the cell. In contrast, genomically integrated assays target the genome directly in its native environment, for example by altering the epigenetic landscape near a candidate regulatory element (e.g. CRISPR interference screens that use dCas9 with an attached KRAB repressor domain [25]). Notably, only a subset of regulatory element predictions based on epigenomic data typically validate in functional characterization assays [3].

While functional characterization assays provide a more direct assessment of regulatory activity, they can also have some limitations. One limitation is their more limited availability across cell types compared to chromatin mark datasets. This is due, in part, to cost and resource constraints associated with these highly specialized assays, as well as technical challenges such as achieving sufficient plasmid transfection efficiency in specific cell types for plasmid based assays [26]. Another drawback for some assays is limited genomic coverage: functional characterization assays often provide readouts for a limited subset of genomic regions, whereas chromatin mark data can be mapped genomewide. Integrative approaches that combine the broad availability of chromatin marks with direct testing of functional assays have the potential to computationally extend the cell type coverage of functional testing assays.

Several existing methods have used data from high-throughput functional characterization assays as training data for supervised methods that predict regulatory activity [27, 28] or predict effects of individual sequence mutations based on features including sequence [29–32]. However, these methods generally focus on scoring sites or bases within the same cell type for which training data is available. As many sequence and transcription factor binding features are cell type specific, a method optimized to make predictions within a cell type it is trained in might be less effective at making predictions that generalize well across cell types. Additionally, the reliance on a single functional characterization assay or dataset, as commonly seen in existing methods, could introduce biases to the predictions, given that technical differences even within the same assay type has been shown to impact the readouts [19].

To address the challenges of predicting regulatory activity genomewide across a range of cell and tissue types, we propose ChromActivity, a computational framework that integrates chromatin marks with a variety of functional characterization datasets. ChromActivity employs a supervised learning approach to generate genomewide regulatory activity predictions and annotations across multiple cell types. ChromActivity is designed to effectively generalize across both cell types and genomic loci and to produce annotations that reflect differences between functional characterization assays. We apply ChromActivity in over one hundred human cell and tissue types to generate a set of genomewide regulatory activity prediction tracks, where each track is based on a model that is specifically trained on one of 11 functional characterization datasets. ChromActivity generates ChromScoreHMM genome annotations, which correspond to combinatorial and spatial patterns in the prediction tracks. ChromActivity also generates ChromScore, a composite genomewide regulatory activity prediction track on a per-cell or tissue type basis that reflects the mean predicted regulatory activity based on the different functional characterization datasets. The ChromActivity framework and associated annotations provides a resource for analyzing gene regulatory activity across a broad range of human cell and tissue types.

## Results

### Overview of the ChromActivity framework

We developed ChromActivity to provide annotations and scores of predicted regulatory activity across human cell and tissue types by leveraging information from both epigenomic data and a variety of functional characterization datasets. ChromActivity does this by first predicting separately for each functional characterization dataset the relative likelihood of each 25-bp genomic interval showing activity based on chromatin mark features across the entire human genome. These individual prediction scores are then integrated to produce ChromScoreHMM annotations, which are unsupervised genome annotations built on top of ChromHMM [4, 13, 14]. The scores are also integrated into a single combined score, ChromScore (Figure 1A, Supplementary Figure S1).

**Figure 1:**
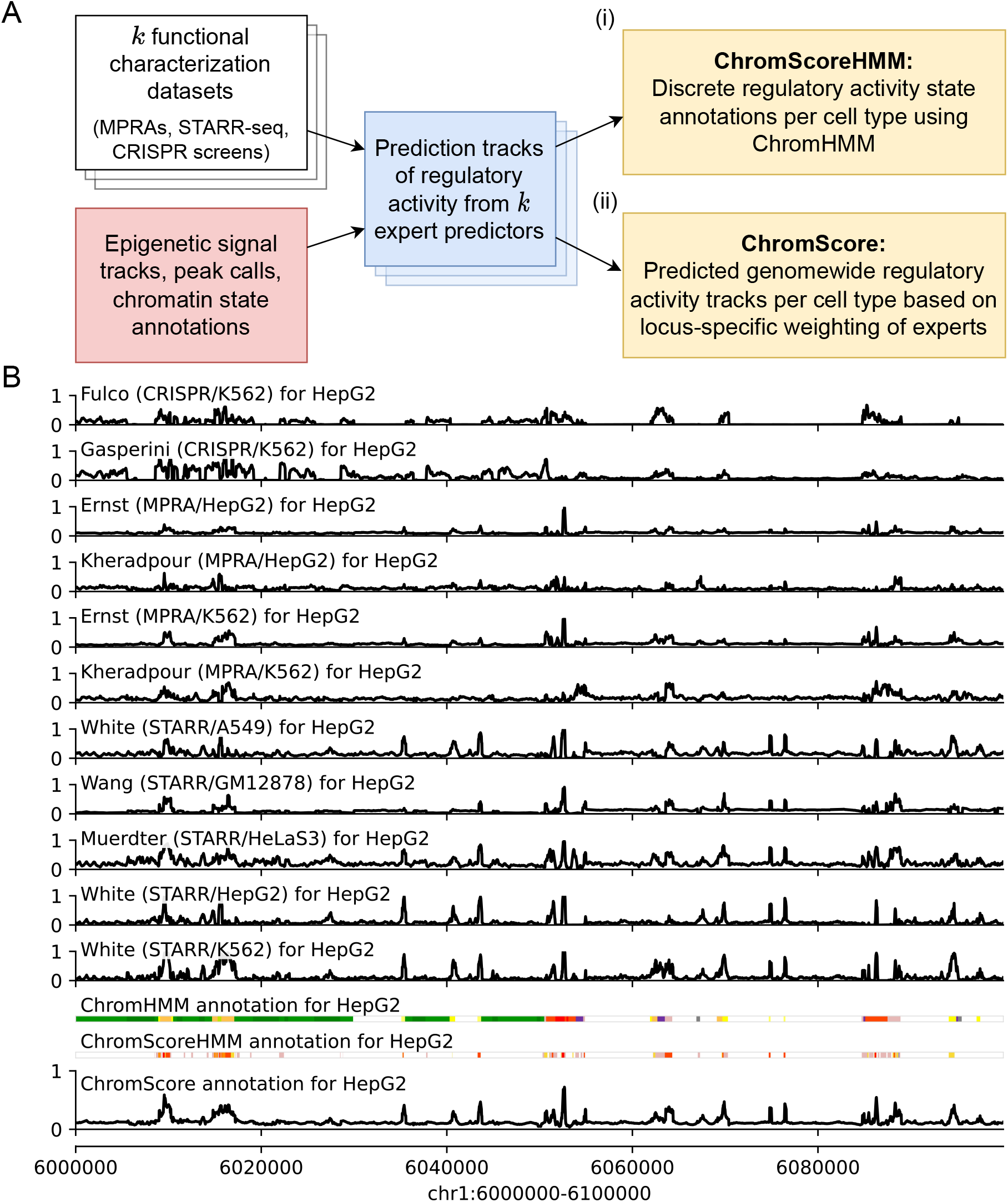
Overview of the ChromActivity framework. **(A)** Flowchart of the ChromActivity framework. ChromActivity takes as input regulatory activity labels from targeted genomic regions from *k* different functional characterization datasets (stacked white blocks, upper left). Using features based on chromatin mark signals, peak calls and chromatin state annotations for the targeted regions (red block, lower left), it trains a separate classifier (“expert”) for each functional characterization dataset. Each expert provides a predicted genomewide regulatory activity score track specific to a functional characterization dataset (stacked blue blocks). ChromActivity then uses the score tracks to generate two complementary outputs reflecting predictions of regulatory activity for each cell type (yellow blocks, right): **(i)** ChromScoreHMM annotations, which are annotations of the genome into states generated by integrating combinatorial and spatial patterns in the expert prediction score tracks using ChromHMM and **(ii)** ChromScore tracks, which are continuous genomewide regulatory activity score tracks based on the mean individual expert scores at each 25-bp interval. See Supplementary Figure S1 for a detailed schematic. **(B)** Visualization of regulatory activity score tracks for each expert, ChromHMM chromatin state annotations (25-state imputed model), the ChromScore track and ChromScoreHMM annotations in HepG2 for genomic interval chr1:6,000,000-6,100,000 (hg19). ChromScoreHMM and ChromHMM color legends are shown in Supplementary Figure S2.

ChromActivity makes predictions for any cell type with chromatin mark data available (For ease of presentation we use the term “cell type” to refer to cell types, tissue types and reference epigenomes collectively). Notably, ChromActivity operates without assuming any functional characterization data is available in the cell types for which it predicts. This is important as currently there are a large number of cell types that have extensive chromatin mark data available, but do not have data from functional characterization assays available.

ChromActivity’s approach is based on the observation that the same chromatin mark patterns generally mark regulatory regions in different cell types, though the location of those patterns can vary between cell types [4, 9]. This contrasts with specific DNA sequence or transcription factor binding patterns which can mark regulatory regions only in specific cell types, and thus we do not use them as features in ChromActivity.

Initially, ChromActivity trains a separate bagging ensemble of regularized logistic regression models for each input functional characterization dataset. These models are trained with labels derived from the readouts for the genomic regions tested by the functional characterization datasets, which include both plasmid-based (MPRAs, STARR-seq screens) [33–37] and genome-integrated assays (CRISPR-dCas9 screens) [25, 38] from multiple different conditions and cell types (Methods). ChromActivity uses features derived from signal and peak calls of individual chromatin marks, as well as chromatin state annotations [13, 14]. In addition to using the signal directly at the tested loci, ChromActivity incorporates spatial information from the signal track by extracting the signal at 25 bp resolution within 2 kb windows centered around the tested loci and then uses principal component analysis (PCA) to reduce the number of these additional signal features per mark from 81 to 3 (Methods). ChromActivity trains each logistic regression ensemble on a single functional characterization dataset, which we term ChromActivity ‘experts’. ChromActivity then applies these experts to make predictions across the entire genome in a large number of cell types, only one of which each expert would have seen training data from.

As individual expert predictions can in some cases disagree on predictions of regulatory activity, ChromActivity uses the individual expert predictions to generate genome annotations corresponding to combinatorial and spatial patterns of top predicted positions of regulatory activity from the different experts within a cell type. To do this, ChromActivity applies ChromHMM [13, 14] with input based on the different expert predictions to generate what we term ChromScoreHMM genome annotations (Methods). Relative to ChromHMM annotations, which are defined directly based on chromatin marks, these states are intended to more directly correspond to regions which have chromatin mark annotations predictive of regulatory activity in all or specific subsets of functional characterization datasets.

In addition, ChromActivity averages predictions from different experts to generate ChromScore, a single genomewide regulatory activity potential score for each cell type. ChromScore provides a numeric score between 0 and 1 of predicted regulatory activity potential for any 25 bp segment of the genome.

### Training and evaluation of ChromActivity experts

We applied ChromActivity to imputed signal tracks and peak calls for ten histone modifications: H3K27ac, H3K27me3, H3K36me3, H3K4me1, H3K4me2, H3K4me3, H3K79me2, H3K9ac, H3K9me3, H4K20me1, histone variant H2A.Z and the DNase-I hypersensitivity (DNase) signal for 127 cell types from the Roadmap Epigenomics compendium [9, 39]. We used imputed data as it enabled us to apply our method with more marks across more cell types in a uniform manner than with observed data. We also included features based on a one-hot encoding of the 25-state ChromHMM chromatin state annotation that was previously trained on the same 12 imputed marks as our features.

We generated binary “activating” and “neutral” labels (Methods) for each genomic locus in 11 functional characterization datasets (Supplementary Table 1). Five different cell types (A549 lung carcinoma, GM12878 lymphoblastoid, HeLa-S3 cervical carcinoma, HepG2 liver carcinoma and K562 myelogenous leukemia cell types) were represented in the functional characterization datasets. Among the 11 datasets, two were CRISPR-dCas9-based assays (Fulco/K562 [38], Gasperini/K562 [25]). Additionally there were nine plasmid-based assays (Methods), which we further classified into four MPRAs (Ernst/HepG2, Ernst/K562 [34], Kheradpour/HepG2, Kheradpour/K562 [33] five STARR-seq-derived assays (Muerdter/HeLaS3 [35], Wang/GM12878 [36], White/A549 [7], White/HepG2, and White/K562 [37]).

The total number of genomic loci used in training each individual expert ranged from 816 to 38,452 (Supplementary Figure S3C). On average, 8.98% of genomic loci in a given dataset was within 100 bp of any locus in any other dataset. Across dataset pairs this overlap varied from 0.01% to complete overlap in the cases of Ernst/HepG2 with Ernst/K562 and Kheradpour/HepG2 with Kheradpour/K562. The fraction of DNase-I hypersensitive sites (Supplementary Figure S3B) and the chromatin state distributions of the loci (Supplementary Figure S3C) also varied across the datasets, which was expected as the datasets employed diverse strategies for selecting loci for testing.

While our main focus is cell type generalization, to establish reference points for the prediction difficulty for each dataset, we first evaluated predictive performance of the experts at distinguishing activating vs. neutral labeled loci in unseen partitions of the same dataset in which they were trained (Supplementary Figure S4A). We found median out-of-sample prediction AUROCs ranged from 0.65 (Kheradpour/K562) to 0.93 (White/HepG2), with mean AUROC across all experts of 0.80. Expert predictive performance generally increased with the number of loci used in training (Spearman correlation: 0.75, Supplementary Figure S4B). The expert models trained on the STARR-seq datasets Muerdter/HeLaS3, Wang/GM12878, White/A549, White/K562, White/HepG2 all had relatively high median AUROCs (0.80, 0.83, 0.89, 0.91 and 0.93 respectively) compared to experts trained on other assay types (average AUROCs of 0.75 for MPRAs, 0.72 for CRISPR-based screens). In addition to the larger size of their training data, another possible contribution to the higher predictive performance of STARR-seq based experts could be explained both by the larger training data sizes and differences in the distribution of loci tested, which in the STARR-seq data include a broader and more diverse set of loci (Methods).

### Genomewide expert predictions

For each of the 127 cell types, ChromActivity computed a score track for each expert predictor reflecting its genomewide regulatory activity predictions (Figure 1B). We quantified the agreement among the individual expert regulatory activity scores based on the mean of pairwise Pearson correlations computed across the genome (Figure 2A, Methods). The different expert predictions exhibited moderate agreement with an average pairwise Pearson correlation of 0.37 across all pairs of 11 score tracks and cell types. The pairwise correlations of experts ranged from -0.14 to 0.90, with the extremes corresponding to experts trained on the pair Gasperini/K562 (CRISPR-based) and Wang/GM12878 (STARR-seq) and the pair White/A549 (STARR-seq) and White/HepG2 (STARR-seq), respectively. We observed higher correlations within predictions from experts trained on plasmid-based (mean correlation 0.51) and CRISPR based (correlation 0.52) functional characterization datasets than the correlations between plasmid-based and CRISPR-based experts (mean correlation 0.09).

**Figure 2:**
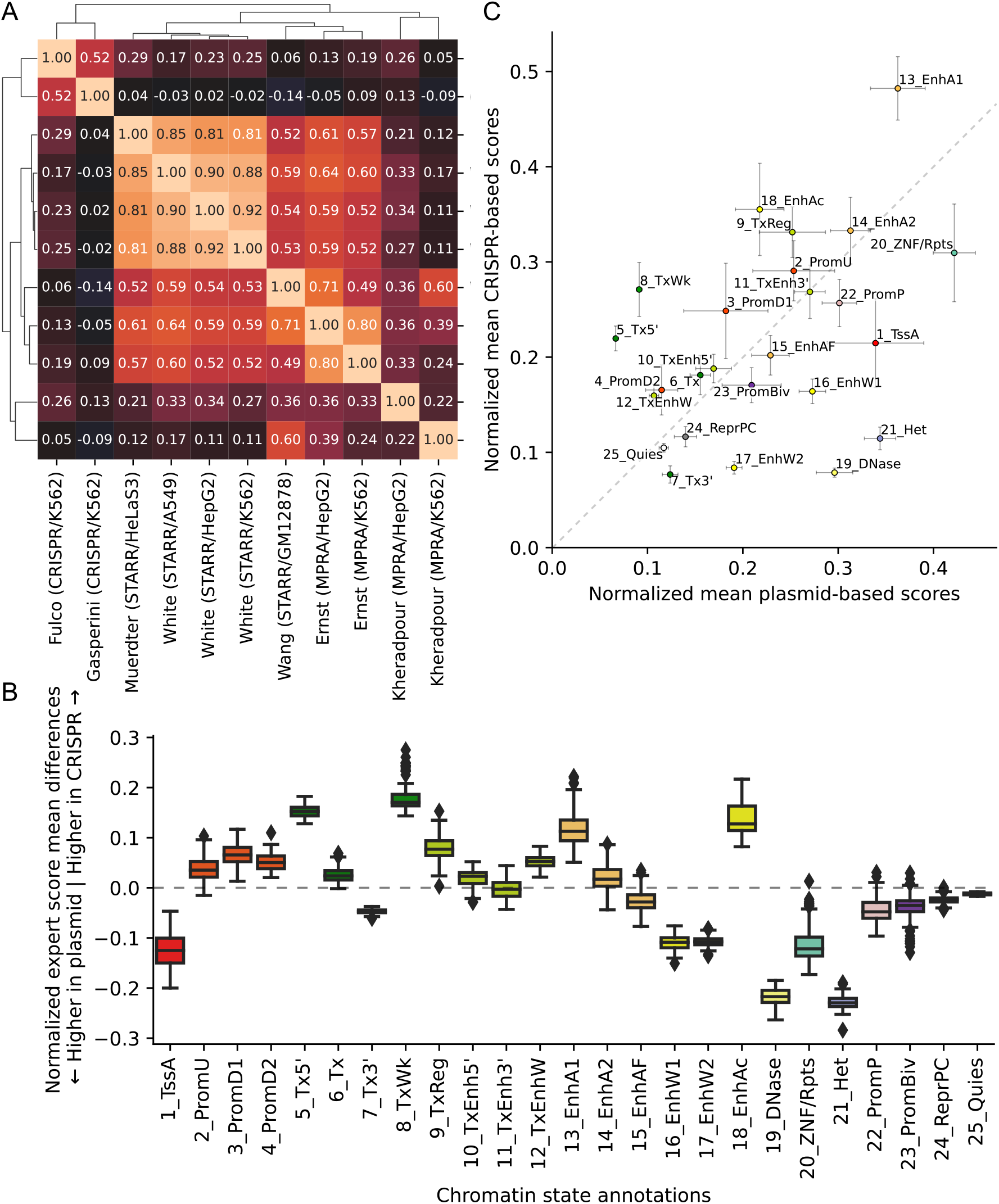
Correlation of individual expert scores and comparison of plasmid-based and CRISPR-based experts. **(A)** Heatmap of mean genomewide Pearson correlations between expert model tracks clustered with hierarchical clustering, averaged over cell types. **(B)** Box plots of mean normalized score differences across cell types between experts trained on 9 plasmid based and 2 CRISPR-based functional characterization datasets in different ChromHMM chromatin states [13, 39]. The boxes represent quartiles and whiskers indicate maximum and minimum score differences between plasmid-based and CRISPR-based experts. Individual mean scores averaged across cell types, for each expert separately, is shown in Supplementary Figure S5. The corresponding box plot distributions of means across cell types for each expert and each state is shown in Supplementary Figure S6. **(C)** Scatter plot of mean normalized expert scores for plasmid-based vs. CRISPR-based functional characterization datasets per chromatin state, averaged over cell types. Error bars indicate standard deviation of score means across cell types.

Correspondingly, clustering the experts based on pairwise correlations of genomewide predictions revealed two main clusters (Figure 2A). The first cluster included predictions from the two CRISPR-based experts, Fulco/K562 and Gasperini/K562. The second cluster included predictions from all but two plasmid experts (average pairwise Pearson correlation 0.67) and itself contained two subclusters. One subcluster included predictions from the three White lab experts and Muerdter/HeLaS3 (average correlation 0.86) and the other contained the Ernst/K562, Ernst/HepG2, and Wang/GM12878 experts (average correlation 0.67). Outside of the two main clusters, there were two experts, Kheradpour/K562 and Kheradpour/HepG2, which had low correlations with each other (0.22) and with predictions from other experts (average correlations 0.28 and 0.19).

We observed considerable variability in the chromatin states prioritized by different experts (Supplementary Figure S5), notably between plasmid-based and CRISPR-based experts. For instance, regions overlapping the heterochromatin-associated 21_Het state had substantially greater normalized predicted activity based on the plasmid-based experts compared to CRISPR-based experts (Figure 2B). This is consistent with DNA sequences that are active in the plasmid context, but are repressed by H3K9me3 marked heterochromatin in the native chromatin context.

### ChromScoreHMM genome annotations

To better understand the relationships between ChromActivity’s expert model predictions and to generate an integrated genome annotation based on them, we developed ChromScoreHMM. ChromScoreHMM identifies combinatorial and spatial patterns within the expert predictions and annotates the genome at 25-bp resolution based on them. ChromScoreHMM starts by binarizing the expert model predictions based on a top 2% threshold computed separately for each expert in each cell type (Methods). ChromScoreHMM then uses the binarized predictions across the cell types as input to ChromHMM [4, 13, 14] to learn a multivariate hidden Markov model. ChromScoreHMM learns a model across cell types using the ‘concatenated’ approach, leading to a shared set of states across cell types but cell type specific assignments. The states capture distinct combinatorial and spatial patterns of expert predictions, and the resulting genome annotation is termed the ChromScoreHMM annotations.

We focused our analysis on a ChromScoreHMM model with 15 states (Methods). We numbered the states in decreasing order of mean emission parameter values (Figure 3A) and divided the states into three subgroups consisting of what we characterized as multi-expert states (States 1-10), single expert states (States 11-14), and the no expert state (State 15) (Supplementary Figure S2). The multi-expert states all had at least two experts with emission probabilities ≥0.20. The single expert states all had a single expert with an emission probability of ≥0.90 and no other experts with emission probability ≥0.10. In the no expert state, all experts had <0.001 emission probability. The multi-expert and single expert states covered in total 5.2% and 3.9% of the genome respectively, while the no expert state was by far the most common state covering 90.8% of the genome.

**Figure 3:**
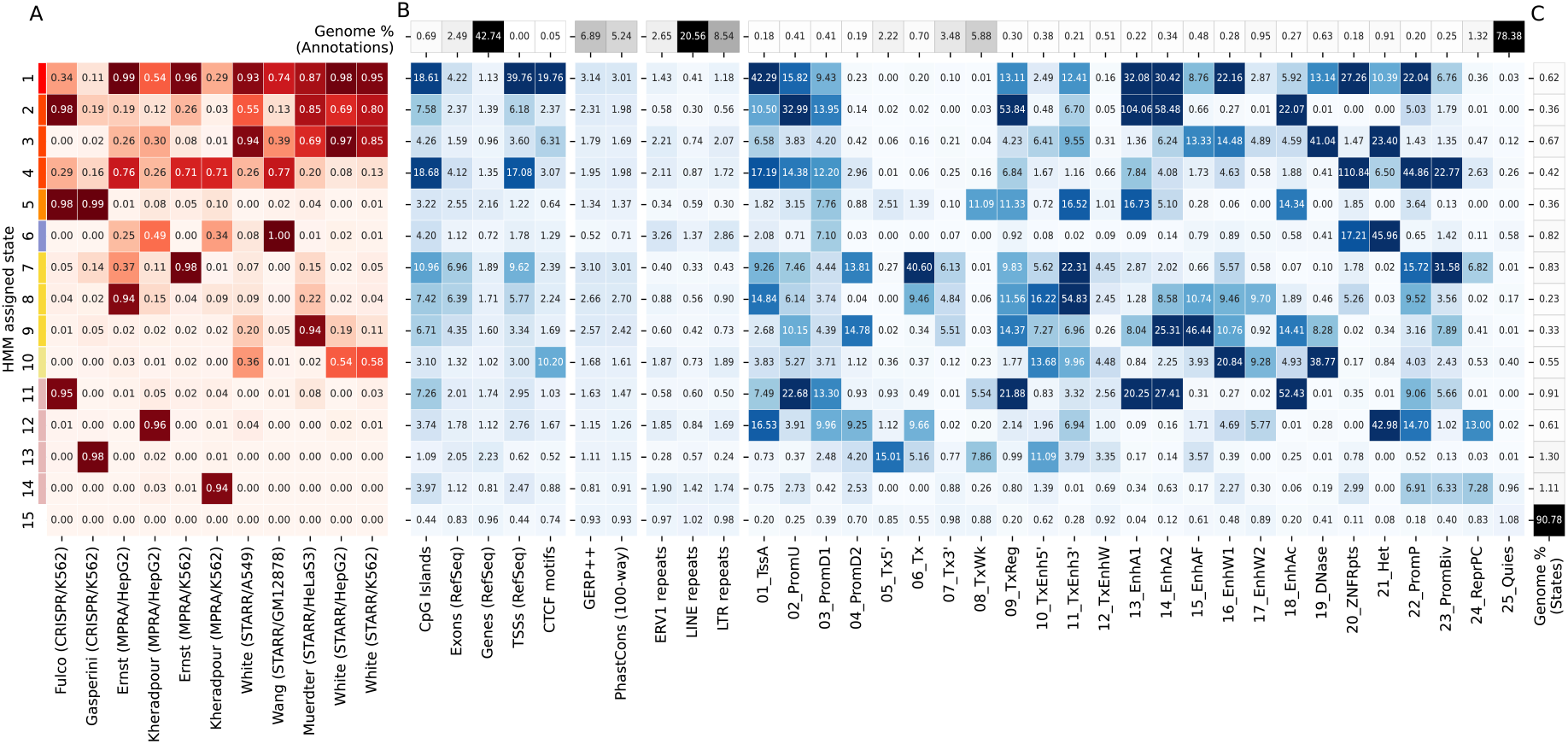
ChromScoreHMM emission parameters and enrichments. **(A)** Emission parameters of a ChromScoreHMM model learned based on combinatorial and spatial patterns of top scoring predictions of each expert (top 2% of predictions, Methods). Each row of the heatmap corresponds to a ChromScoreHMM state (states 1-15, color legend on left margin) and each column a different input expert model. Emission parameter values correspond to the probability in that state of observing a top scoring prediction for that expert model. **(B)** Overlap fold enrichments for **(1)** sequence and gene annotations: CpG islands [40], exons, gene bodies and transcription start sites from RefSeq [41], CTCF motifs from HOMER [42], **(2)** evolutionary conservation related annotations: GERP++ [43], PhastCons 100 vertebrates conservation [44], **(3)** ERV1, LINE and LTR repeat elements from RepeatMasker [45], **(4)** ChromHMM annotations, 25-state model [13, 39]. ***Top row:*** Percentage of the genome occupied by the annotation. **(C)** Percentage of the genome assigned to each ChromScoreHMM state. See Supplementary Figures S7 and S8 for additional enrichments. Red shading: emission parameters, blue shading: fold enrichments, black shading: genome percentages. Enrichments and percentages are medians across cell types.

Among the multi-expert states, State 1 was the only state that had emission probabilities >0.10 for all 11 experts. It had relatively high emission probabilities (>0.50) for eight of the experts based on MPRA and STARR-seq datasets, and moderate emission probability (0.10-0.50) for the two experts based on CRISPR-dCas9 datasets and one MPRA dataset. State 2 had high emission probabilities for the Fulco/K562 (CRISPR-based) expert and all but one STARR-seq based expert. State 3 had moderate or high emission probabilities for all the STARR-seq based experts and two of the MPRA experts. State 4 had moderate or high emission probabilities for all experts with the exception of White/HepG2. In contrast, State 5 was dominated by two CRISPR-based experts, with emission probabilities >0.98 for both, while highest non-CRISPR expert emission probability was 0.10. States 6-9 had one expert that had a very high emission probability (≥0.95) and moderate emissions for one or two other experts. For instance, State 7 had very high (0.98) and moderate (0.37) emission probabilities for the Ernst/K562 and Ernst/HepG2 experts, respectively, while State 8 had high (0.94) and moderate (0.22) emission probabilities to the Ernst/HepG2 and Muerdter/HeLaS3 experts, respectively. State 10 was uniquely associated with the three White lab STARR-seq datasets, with each associated expert having a moderate emission probability and other experts having low emission probability (<0.05), suggesting that this state may be capturing aspects of this particular STARR-seq protocol or other types of batch effects.

The four single expert states (States 11-14) were each associated with one expert. States 11 and 13 were associated with the CRISPR-based experts Fulco/K562 and Gasperini/K562 respectively, while States 12 and 14 were associated with the experts for Kheradpour HepG2 and K562 datasets, respectively. The experts associated with the single expert states had below average pairwise score correlations with other experts (0.21 and 0.04 for Fulco/K562 and Gasperini/K562, respectively, and 0.19 and 0.28 for Kheradpour/K562 and Kheradpour/HepG2, respectively, compared to average pairwise correlation over all pairs of 0.37, Figure 2A). These results suggest that these single expert states might be capturing dataset-specific signals or biases.

### Enrichment analysis of ChromScoreHMM states

To better understand genomic properties of individual ChromScoreHMM states, we computed state enrichments for various genomic annotations. Some of the annotations used for the enrichments were also used as input to ChromActivity, such as ChromHMM chromatin states (Figure 3) and chromatin mark peak calls (Supplementary Figure S8C). Other annotations were independent of ChromActivity’s predictions, including CpG islands, CCCTC-binding factor (CTCF) motifs, transcription start sites, exons, gene bodies, various repeat elements and evolutionarily conserved elements (Figure 3). We also computed the proportion and fold enrichments of the ChromScoreHMM states in the genomic neighborhood of TSSs (Supplementary Figure S10). Additionally, we computed the normalized average prediction score for individual experts in each state (Supplementary Figure S7C).

Seven of the ChromScoreHMM states showed strong (>10 fold) enrichments for at least one of the active enhancer or flanking chromatin states 13_EnhA1, 14_EnhA2 or 15_EnhAF (Figure 3), including 6 multi-expert (States 1, 2, 3, 5, 8, 9) and one single expert state (State 11, corresponding to the Fulco/K562 expert). State 2 (most associated with Fulco/K562, Muerdter/HeLaS3 and the White STARR-seq experts) had the strongest enrichments for the active enhancer states 13_EnhA1 and 14_EnhA2 among all the ChromScoreHMM states, with median fold enrichments of 104.0 and 58.5 fold respectively, while State 9 had the highest enrichment for the active enhancer flanking state, 15_EnhAF (46.4 fold).

Among all the states, State 1 (associated with broad expert activity) was most strongly enriched for both the TSS associated chromatin state 1_TssA (42.3 fold) and annotated TSSs themselves (39.8 fold), with a sharp peak in fold enrichment just around the TSS that levels off to approximately 1.7 fold 2 kb upstream and downstream of the TSS (Supplementary Figure S10C). State 1 was also highly enriched for CpG islands (18.6 fold) and CTCF motifs (19.8 fold).

Of the eight states that did not show strong enrichment for any of the active enhancer or flanking states, three of them still showed moderate enrichment for conserved bases (1.3 - 3.0 fold) including two multi-expert states (States 7, 10) and one single expert state (State 12). State 4 (associated with moderate to high enrichment in many experts) was also notable in that it was strongly enriched for CpG islands (18.7 fold) and for the chromatin states associated with poised promoters (22_PromP, 44.9 fold) and ZNF genes and repeats (20_ZNF/Rpts, 110.8 fold). State 7 (most associated with Ernst/K562 and Ernst/HepG2) was notable in that it showed the strongest enrichment for the bivalent promoter state (23_PromBiv, 31.6 fold) and also showed strong enrichment (>10 fold) for two transcribed enhancer states (6_Tx and 11_TxEnh3’). State 10, which was associated with a subset of the STARR-seq-based experts (White/A549, White/HepG2, White/K562), showed strong enrichment for a DNase specific chromatin state (19_DNase, enrichment 38.8 fold). The presence of DNase without histone modifications is often associated with CTCF binding and candidate insulator regions [46]. Consistent with that, State 10 had a 10.2 fold enrichment for CTCF motifs.

Interestingly, some ChromScoreHMM states showed enrichment for both chromatin states associated with repression and activation. For instance, State 12 (the single expert Kheradpour/HepG2 state) was strongly enriched for the polycomb repressed chromatin state (24_ReprPC, 13.0 fold), the repressive heterochromatin associated chromatin state (21_Het, 43.0 fold) and the poised promoter state (22_PromP, 14.7 fold), but also the active TSS state (1_TssA, 16.5 fold). Similarly, State 3 was enriched for the repressive 21_Het state (23.4 fold) while also being enriched for moderately active states like 15_EnhAF (13.3 fold) and 16_EnhW1 (14.5 fold).

Three states (States 6, 14 and 15) were predominantly associated with repressive or quiescent genomic chromatin states. State 6 (most associated with Wang/GM12878, Kheradpour/HepG2 and Kheradpour/K562), had the strongest enrichment of any state for the 21_Het chromatin state (46.0 fold) and also for LTRs (2.9 fold) while having the strongest depletion of conserved bases (0.71 fold). State 14 (the single expert state for Kheradpour/K562) was enriched for the repressive poised promoter (22_PromP), bivalent promoter (23_PromBiv) and repressed polycomb (24_ReprPC) states (6.9, 6.3 and 7.3 fold respectively). Additionally, State 14 showed the weakest depletion for the Quiescent chromatin state (25_Quies) among single or multi expert states (0.96 fold) with the Quiescent chromatin state comprising 75% of the state. State 15 (the no expert state) was the only ChromScoreHMM state to show enrichment for the Quiescent chromatin state (1.1 fold).

Notably, all ChromScoreHMM states with high emission parameter values for CRISPR-based experts (States 2, 5, 11, 13) were depleted for the 21_Het heterochromatin chromatin state (Figure 3), which was not the case in general for states with high emission parameter values for plasmid-based experts. This is consistent with our analysis of individual CRISPR-based and plasmid-based experts (Figure 2B), which showed 21_Het was being assigned higher scores by the plasmid-based experts compared to the CRISPR-based experts, likely marking regulatory sequences that are repressed in their native chromatin context.

ChromScoreHMM annotations displayed substantial variation in mean gene expression between states and specific positions relative to the TSS (Supplementary Figure S11). Most ChromScoreHMM states were more enriched at TSSs of high expression genes than low expression genes (Supplementary Figure S12). States 14 and 15, which were among the states associated with repressive or quiescent genomic regions, were exceptions to this. State 6, which was also predominantly associated with repressive or quiescent chromatin states, was more enriched for low expression genes upstream and downstream of the TSS, but was more enriched for high expression genes at the TSS (Supplementary Figure S12B). Meanwhile, States 5 and 13, which are mainly associated with CRISPR-based experts, were more enriched downstream of the TSSs of high expression genes compared to low expression genes.

### ChromScore regulatory activity predictions

ChromActivity also averages the outputs of its individual expert predictions to generate a cell type specific regulatory activity score, termed ChromScore (Figure 4A, Methods). ChromScore provides a single continuous score track for each cell type, where higher scores correspond to higher average predicted regulatory activity potential (Figure 4A).

**Figure 4:**
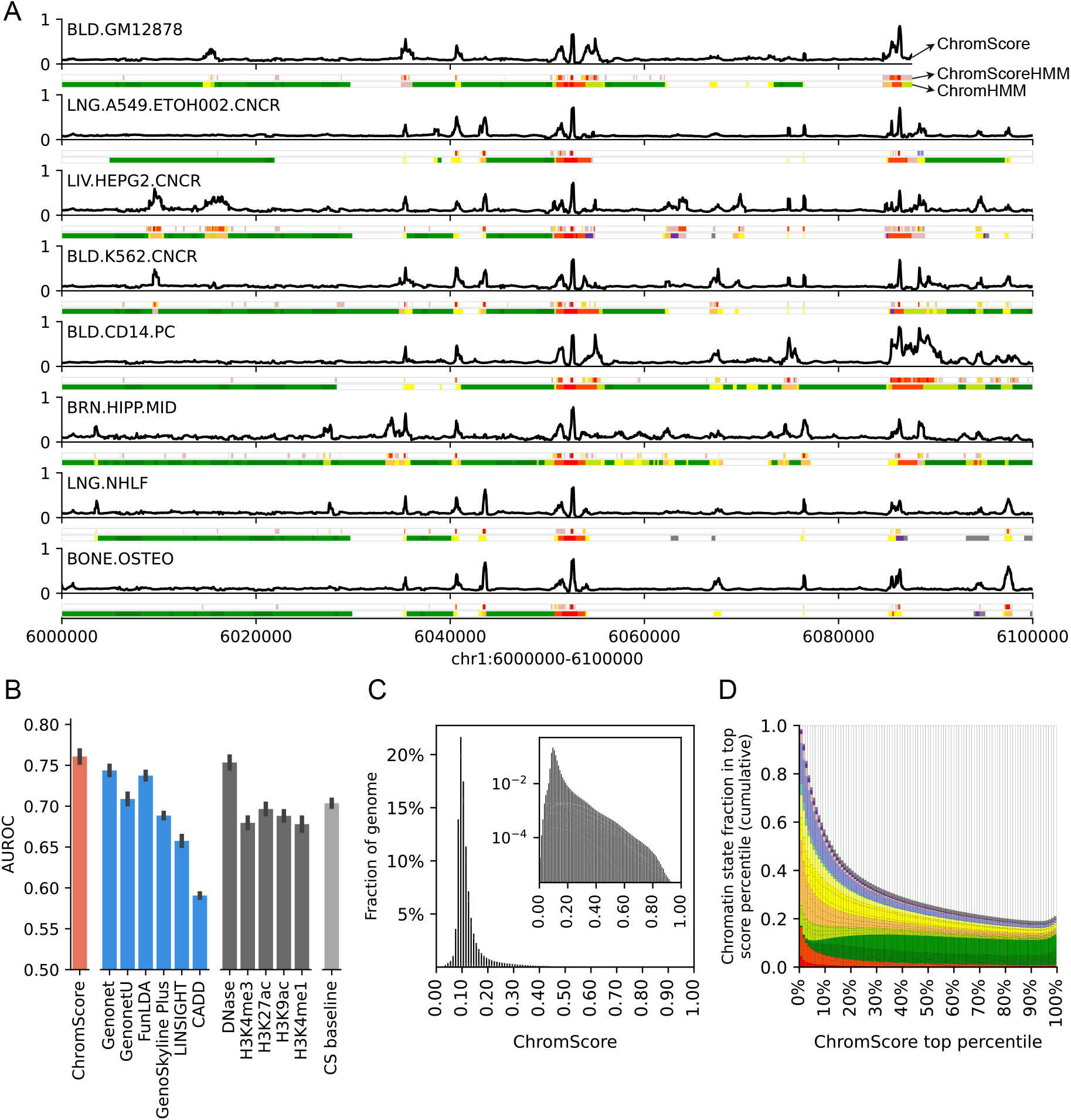
ChromScore tracks, cell type generalization performance evaluations and score distributions. **(A)** Visualization of ChromScore tracks in eight cell types shown above ChromScoreHMM and ChromHMM annotations in the same cell types for genomic interval chr1:6,000,000-6,100,000 (hg19). The cell types shown represent examples of both those with and without functional characterization training data (cell types with training data: GM12878, A549, HepG2, and K562; cell types without training data: CD14 primary monocytes, brain hippocampus cells, NHLF lung fibroblast primary cells, and osteoblast primary cells). **(B)** A comparison of cell type generalization performance of ChromScore to existing scores, single marks, and a chromatin state baseline. The bars correspond to the mean area under receiver operator characteristic (AUROC) across 11 functional characterization datasets. The first bar shows the performance of ChromScore. For ChromScore evaluations, expert models trained on the same cell type as the evaluation dataset were not used. The next six bars show the performance of existing scores [27, 47–51], which are followed by bars for the imputed signal tracks for DNase I hypersensitivity, H3K4me3, H3K27ac, H3K9ac, and H3K4me1. The last bar shows the mean ensemble of the chromatin state baseline models for all datasets (CS baseline, Methods). Error bars indicate standard error across evaluations. **(C)** Genomewide distribution of ChromScore values, averaged over cell types. Inset: log scaled. **(D)** Cumulative chromatin state fraction for top ChromScore percentiles. Each bin corresponds to an additional top 1% of scores. See Supplementary Figure S2 for chromatin state color legend.

We investigated if ChromScore, which was trained based on functional characterization assay data in a limited number of cell types, would generalize to new cell types without functional characterization data. To evaluate the cell type generalization performance of ChromScore to predict regulatory activity in unseen cell types, first we generated modified versions of our ensemble models in which functional characterization datasets of each cell type were removed from the training data. Next, we generated and evaluated ChromScore tracks for the held out cell types using the modified models at loci not seen in training (Figure 4B, Supplementary Figure S13).

We compared the ChromScore predictions to a set of baselines and existing score tracks from other methods for predicting activating vs. neutral labels of loci tested with functional characterization assays. The baselines included those based on individual chromatin marks and one based on chromatin state assignments (Methods). The existing score tracks included several scores that provided cell type specific regulatory activity estimates integrating multiple epigenomic datasets (GenoNet, GenoNet-U [27], FunLDA [47], Genoskyline Plus [48]). In addition, we compared to two scores that also integrate epigenomic annotations, but do so in a non-cell type specific manner and also consider a diverse set of other annotations, CADD [49] and LINSIGHT [50] (Methods).

ChromScore predictions had a substantially higher mean AUROC score (0.76) relative to all the baselines (AUROC range 0.67-0.70) except relative to DNase signal in which it was marginally better (0.75) (Figure 4B). Among the existing scores we evaluated, AUROCs ranged from 0.59 (CADD) to 0.74 (Genonet and FunLDA). Comparing ChromScore’s predictive performance to its underlying experts indicated that ChromScore performed similar to or better than the highest scoring experts in the majority of evaluations (Supplementary Figure S14). ChromScore performed better in plasmid-based dataset evaluations (mean AUROC 0.79) compared to CRISPR-based dataset evaluations (mean AUROC 0.59, Supplementary Figure S15), possibly because it was trained on more plasmid-based datasets. Notably, while ChromScore and DNase signal showed similar cell type generalization performances, they were only moderately correlated across cell types (median Spearman correlation 0.26, Supplementary Figure S16), and the chromatin state distributions of top ChromScore regions differed considerably from top DNase regions (Supplementary Figure S17).

The median ChromScore across the genome and all 127 cell types was 0.10 (Figure 4C), with top-scoring genomic regions (highest 2% genomewide) having an average ChromScore > 0.35. A number of the chromatin states with high mean ChromScores (Figure 5C) and high fold enrichments within top-scoring genomic regions (Supplementary Figure S17A) across cell types included states typically associated with regulatory activity, such as 13_EnhA1 (mean score 0.41, fold enrichment 29.40 fold), 14_EnhA2 (mean score 0.35, fold enrichment 23.43) and 1_TssA (mean score 0.32, fold enrichment 17.46). Interestingly, other chromatin states such as 20_ZNF/Rpts (mean score 0.41, 18.01 fold) and 21_Het (mean score 0.32, 14.34 fold) also displayed high mean scores and top-scoring region enrichments. The high mean scores and top-scoring region enrichments of 20_ZNF/Rpts and 21_Het appeared to be mainly driven by plasmid-based experts which, as previously shown, were more likely to assign higher scores on average to 20_ZNF/Rpts and 21_Het-annotated genomic regions (Supplementary Figure S5, S6).

**Figure 5:**
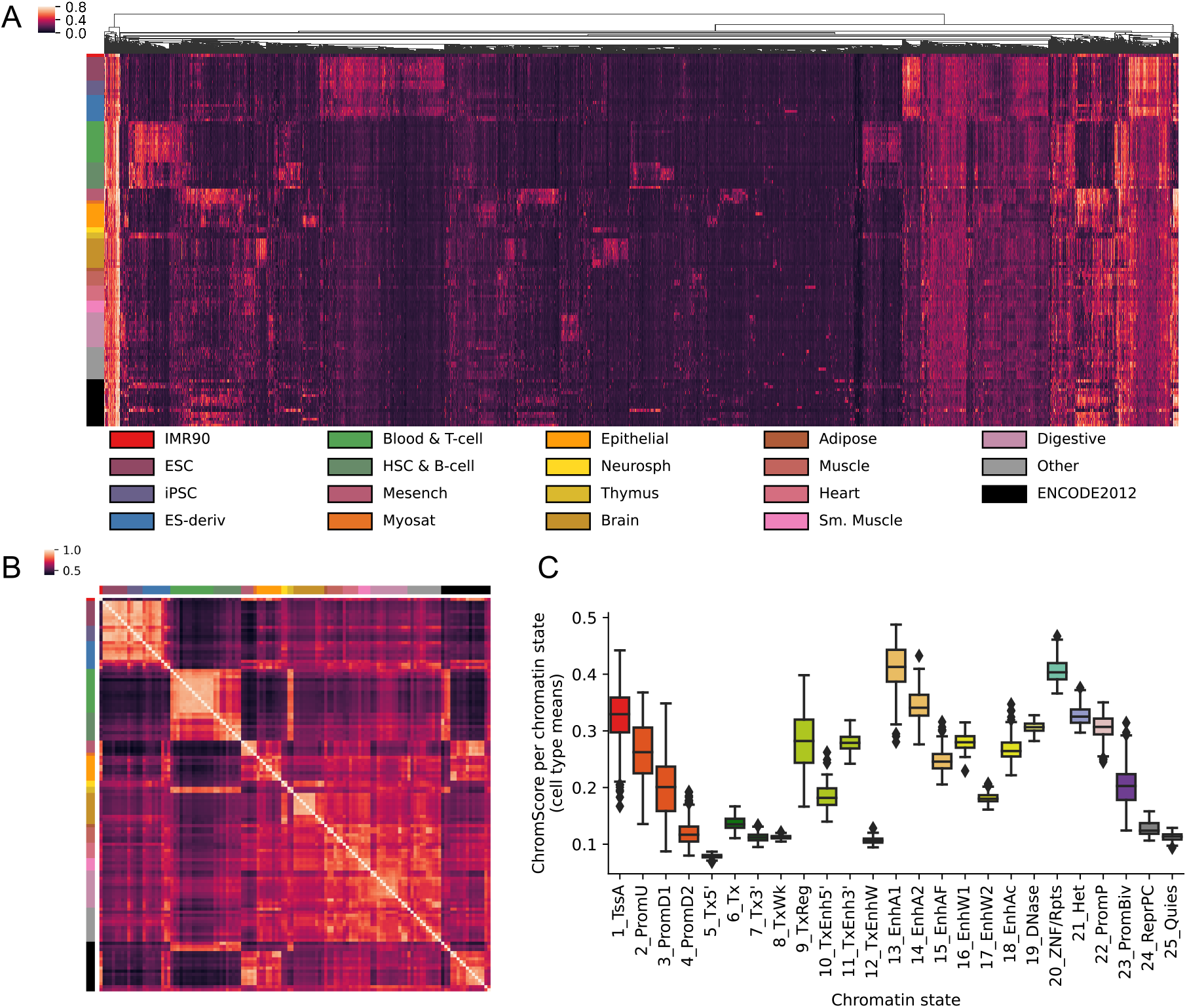
ChromScore across cell types. **(A)** Heatmap showing ChromScores at 20,000 randomly selected bases across the genome (columns) that had a score difference of >0.25 between at least two cell types for 127 Roadmap Epigenomics cell types (rows) (Methods). Columns are hierarchically clustered and rows are sorted based on Roadmap Epigenomics tissue groups [9]. The tissue groups of the rows are indicated on the left and their color legend is displayed at the bottom. **(B)** Heatmap of ChromScore Pearson correlations across all pairs of 127 cell types, which are ordered and colored as in A. **(C)** Distribution of mean ChromScores per chromatin state per cell type.

Analyzing ChromScore across many cell types enabled us to identify some genomic loci that were predicted to show near-universal activity across a diverse range of cell types. Approximately 0.19% of loci across the genome were predicted to be highly active (top 2% ChromScore) in over 90% of all cell types. We also observed that cell types that were more biologically similar had greater correlation in their ChromScore (Figure 5A, B). In particular, cell types within the same Roadmap Epigenomics tissue group [9] had an average Pearson correlation of 0.80 compared to a correlation of 0.62 for predictions crossing different tissue groups, reflecting ChromScore’s ability to capture cell and tissue-specific behavior.

We analyzed fold enrichments for genomic repeat elements in top-scoring ChromScore regions (top 2%, Supplementary Figure S18A,E), and observed enrichments for long terminal repeats (LTRs, fold enrichment 1.71), particularly the endogenous retroviral sequence 1 (ERV1) subclass (fold enrichment 2.04), and depletions for long interspersed nuclear elements (LINEs, fold enrichment 0.56) and short interspersed nuclear elements (SINEs, fold enrichment 0.41). The enrichment for LTRs is consistent with previous reports showing LTRs association with activating gene expression [34, 52–54]. Top DNase regions by signal, in comparison, were depleted for LTRs, LINEs and SINEs (Supplementary Figure S18D,H). Plasmid-based experts and CRISPR-based experts prioritized different repeat classes, with LTRs being enriched in bases prioritized by plasmid-based experts but depleted in CRISPR-based experts and SINEs including the subclass of Alu elements showing the opposite trend (Supplementary Figure S18B,C,F,G). This could suggest LTRs being repressed in the genome but drive expression in a plasmid context. The enrichment of Alus in bases prioritized by CRISPR-based experts is consistent with the enrichment of both for transcribed regions [55] (Supplementary Figure S5).

ChromScore moderately correlated with in-vivo gene expression at the TSS (Pearson correlation 0.41, Supplementary Figure S19A, Methods). However, some chromatin marks showed stronger correlations, such as H3K9ac (Pearson correlation 0.59 at TSS+500 bp). We note that correlations for ChromScore were not necessarily expected to surpass that of all chromatin marks with expression, since ChromScore heavily relies on plasmid-based experts, which while providing an assessment of the inherent regulatory activity of a DNA sequence, do not reflect the full in-vivo chromatin context. Correlation patterns varied across functional characterization assay types and individual experts (Supplementary Figure S19B), with CRISPR-based experts showing higher correlations upstream and downstream of the TSS (Supplementary Figure S19C) but lower correlations at the TSS compared to plasmid-based experts (Mean Pearson correlations 0.32 for plasmid-based experts, 0.23 for CRISPR-based experts, Supplementary Figure S19D). This observation is consistent with the lower CRISPR expert scores observed in the 1_TssA chromatin state, which is primarily associated with active TSSs, compared to those of plasmid-based experts, as well as the higher scores noted in upstream promoter (2_PromU) and downstream promoter (3_PromD1, 4_PromD2) chromatin states (Supplementary Figure S5). These findings highlight the distinct patterns among ChromScore expert tracks in predicting gene expression around the TSS.

## Discussion

We introduced ChromActivity, a computational framework that predicts gene regulatory element activity across diverse cell types by integrating information from chromatin marks and multiple functional characterization datasets. ChromActivity first trains a set of ‘experts’ with each expert trained using a different individual functional characterization dataset. It then applies these trained predictors to make predictions for each cell type. Using these predictions, ChromActivity produces two complementary integrative outputs for each cell type. One of them is ChromScoreHMM, which annotates the genome into states representing combinatorial and spatial patterns in the expert’s regulatory activity track predictions. The other is ChromScore, which is a cell type-specific continuous numerical score of predicted regulatory activity potential across the genome based on combining the individual expert predictions. We applied ChromActivity using chromatin mark data from 127 cell types in the Roadmap Epigenomics compendium and data from 11 functional characterization datasets.

We observed that different experts prioritized different subsets of the genome, in some cases corresponding to the assay or experimental protocol of the functional characterization dataset it was trained on. For example, plasmid-based experts on average assigned higher regulatory activity prediction scores to H3K9me3 heterochromatin-associated genomic regions compared to CRISPR-based experts, which was expected as the plasmid based experts tested loci outside of their native chromatin context (Figure 2B). These differences allowed us to distinguish genomic regions with likely H3K9me3-associated repressive activity from inactive regions. We also observed differences between CRISPR-based and plasmid-based experts in terms of their correlations with gene expression at and around the TSS and their predictions of regulatory activity for different classes of repeat elements. Given these differences, specific applications may benefit from utilizing either plasmid-based or CRISPR-based expert predictions, or different ChromScoreHMM states. For example, plasmid-based expert tracks and associated ChromScoreHMM states could be preferred for applications focused on predicted regulatory activity inherent in genomic sequences, independent of any regulatory effect of chromatin marks, while the CRISPR-based expert tracks and associated ChromScoreHMM states could be preferred for applications focused on predicted regulatory activity in the native genomic context.

Some of the ChromScoreHMM states corresponded to genomic regions with predicted regulatory activity in different types of functional characterization assays, while others were more specific to a specific assay or likely associated with dataset-specific signals or biases. We showed that ChromScoreHMM states corresponded to substantial enrichment differences for various annotations, including gene annotations, repeat elements, chromatin states and chromatin mark peaks. Further, the spatial distribution of ChromScoreHMM states relative to the TSSs of nearby genes varied depending on the expression of the genes. As expected, most states were more enriched at or around the TSSs of high expression genes compared to low expression genes, except for the few states associated with repressive or quiescent genomic regions.

ChromScoreHMM, while building on the ChromHMM method, provides a distinct genome annotation that complements ChromHMM annotations. In particular, ChromScoreHMM annotations are defined based on combinatorial and spatial patterns in supervised predictions of regulatory activity corresponding to different functional characterization datasets, while ChromHMM annotations are defined directly based on the combinatorial and spatial patterns of chromatin marks. ChromScoreHMM annotations thus more directly correspond to different classes of predicted regulatory activity, while ChromHMM annotations can capture chromatin mark patterns not expected to correspond to differences in regulatory activity reflected in functional characterization assays. In particular, high emission parameters for a state in a ChromScoreHMM model can be directly interpreted to be associated with high predicted regulatory activity based on one or more functional characterization datasets, which is not the case for ChromHMM models.

ChromScore is based on an ensemble of predictors trained on a variety of functional characterization datasets thus avoiding an overreliance on the biases associated with any one dataset. We showed the generalizability of ChromScore predictions across cell types through evaluations of predictive performance in unseen cell types. Top ChromScore regions were highly enriched for enhancer chromatin states as well as classes of repeat elements previously shown to be associated with regulatory activation [34, 54]. We also showed that the predictions across 127 cell types exhibited cell type-restricted activity corresponding to known biological groupings of cell types.

There are several potential avenues for future work building on the current ChromActivity framework. One avenue would be to expand the set of functional characterization datasets used as input to ChromActivity, including adding additional recent CRISPR-based ones and datasets from additional cell types. A challenge to incorporating many additional functional characterization datasets in addition to availability has been the lack of uniform processing. However, this is changing with additional uniformly processed datasets beginning to accumulate in repositories (Luo et al., 2020), facilitating their inclusion in future models. A second avenue for future work would be to develop an improved way to combine expert predictions into a score other than the current strategy of averaging of predictions. This could potentially involve an approach that assigns different weights to different experts globally, for instance based on an estimated level of the noise for the labels on which it was trained, or in a locus specific manner based on how similar the locus is to those for which the expert was trained. Another avenue of future work would be to extend ChromActivity to directly predict loci that are repressed. We designed ChromActivity to focus on predicting activation as the information in the functional characterization datasets that we considered for repression was more limited and inconsistent. However some functional characterization datasets are informative of repression [33, 34] and thus could be used in an extended framework that directly considers repression. Future work could investigate applying ChromActivity to additional cell types in human as well as to non human species. However, we expect ChromActivity to already be a resource for analyzing and interpreting the human regulatory genome across diverse cell types.

## Methods

### Dataset selection and label extraction

We derived labeled training data from 11 functional characterization datasets for ChromActivity (Supplementary Table 1). All datasets were of experiments in cell types for which there was matched uniformly processed chromatin mark data available from the Roadmap Epigenomics consortium. The chromatin mark data for these cell types were all originally generated by the ENCODE project consortium [7].

The individual datasets we used in abbreviated notation are: Ernst/HepG2, Ernst/K562 [34], Kheradpour/HepG2, Kheradpour/K562 [33], Muerdter/HeLaS3 [35], Wang/GM12878 [36], White/A549 [7], White/HepG2, White/K562 [37], Fulco/K562 [38], and Gasperini/K562 [25]. The cell types covered by the individual datasets are: A549 lung carcinoma (epigenome identifier E114), GM12878 lymphoblastoid (epigenome identifier E116), HeLa-S3 cervical carcinoma (epigenome identifier E117), HepG2 liver carcinoma (epigenome identifier E118) and K562 myelogenous leukemia (epigenome identifier E123).

ChromActivity treats predicting regulatory activity as captured by functional characterization assays as a binary classification task and specifically focuses on differentiating activating regions from assumed neutral regions. For input into ChromActivity, we defined binary “activating” vs. “neutral” labels for each genomic region in each functional characterization dataset using dataset specific procedures described below. We note that for datasets that had a reported set of repressive sequences (Ernst/HepG2, Ernst/K562, Kheradpour/HepG2, Kheradpour/K562) we decided to exclude them from training the corresponding expert, while for other datasets the neutral sequences may include some repressive sequences. We also provided ChromActivity a “reference nucleotide” within each region used for training, which we selected as the base we considered most likely representative of the regulatory activity. The specific procedure for selecting the base (e.g. center of construct, nucleotide with the highest signal) depended on the functional characterization dataset and are described below.

The Ernst/HepG2 and Ernst/K562 datasets [34] used a dense tiling of MPRA constructs combined with the SHARPR computational method to assign continuous regulatory activity scores to 5 bp intervals within the tiled regions. For each individual tiled region, we identified the position with absolute maximum value and defined S_absmax_ of the region to be the value at that position if non-negative and otherwise the negative of it. We assigned activating labels to tiled regions with S_absmax_ values exceeding 1, and neutral labels to regions with S_absmax_ values between -1 and 1. We filtered any regions with scores under -1 to exclude likely repressive regulatory regions from the training dataset. This procedure yielded 2405 activating and 10,894 neutral regions for Ernst/HepG2 and 2519 activating and 10,162 neutral regions for Ernst/K562. The reference nucleotide for each region was the center base of the 5 bp interval with the highest absolute maximum SHARPR score.

For the MPRA datasets Kheradpour/HepG2 and Kheradpour/K562 [33], we used the precomputed p-values associated with regulatory activity for each construct against scrambled controls. The constructs with regulatory activity under the expressed p-value threshold of 0.05 were labeled activating and the rest were labeled neutral. This yielded 541 activating and 1548 neutral regions for Kheradpour/HepG2 and 347 activating and 1742 neutral regions for Kheradpour/K562. The reference nucleotide for each region was the center nucleotide of the sequence motif originally used in the experimental design, which also was the center nucleotide of the construct. We excluded any synthetic sequences not represented in the genome from the dataset.

For the STARR-seq based datasets White/A549, White/HepG2, White/K562, we obtained STARRPeaker 1.0 [37] peak calls with ENCODE accessions ENCFF646OQS, ENCFF047LDJ and ENCFF045TVA respectively. We assigned activating labels to the top 10% of the peak calls by the normalized signal output/input fold change value. For the neutral regions, we randomly selected bases from the genome, excluding any that overlapped the ENCODE list of excluded regions [56]. For each activating region, we picked three neutral regions from the genome. This procedure yielded 6929 activating and 20,787 neutral regions for White/A549, 5199 activating and 15,597 neutral regions for White/HepG2 and 3571 activating and 10,713 neutral regions for White/K562. The reference nucleotide for each region was the center nucleotide of the peak, which corresponded to the base with the highest normalized signal output/input fold change value.

For Muerdter/HeLaS3 [35], which was also a STARR-seq dataset, we obtained peak calls from https://data.starklab.org/publications/muerdter_boryn_2017/peaks_inhibitor_correctedEnrichment4_supp.table3.tsv, which corresponded to STARR-seq peaks with corrected fold-enrichment values above 4, the same threshold used for peak calling in [35]. We applied the same random regions selection procedure as above to generate three neutral regions for each activating region. This yielded 9613 activating and 28,839 neutral regions for Muerdter/HeLaS3. The reference nucleotide was the center of the peak for each region.

The Wang/GM12878 dataset [36] is based on a combined experimental and computational functional characterization method called High-resolution Dissection of Regulatory Activity (HiDRA). The experimental part of the method is based on a variant of STARR-seq called ATAC-STARR-seq, which first applies a selection step based on ATAC-seq to identify regions of open chromatin and then applies STARR-seq to these selected regions instead of the whole genome. For Wang/GM12878, we first obtained peak calls for “HiDRA driver elements” identified by the HiDRA-SHARPR2 pipeline and the HiDRA RNA/DNA ratio score track (GEO accession GSE104001). We then assigned activating labels to the driver elements with RNA/DNA ratios above 1. To generate the neutral regions, we randomly selected nucleotides from “HiDRA tiled regions” that were not also in “HiDRA active regions”, maintaining a label ratio of three neutral regions for each activating region. This yielded 2409 activating and 7227 neutral regions for Wang/GM12878. The reference nucleotide was the center of the peak for each region.

For Gasperini/K562, a CRISPR-dCas9 dataset [25], we obtained the data for the scaled up experiment from GEO (accession GSE120861), which included genomic regions targeting DNase-I hypersensitive sites with various combinations of H3K27ac, p300, GATA1 and RNA Pol II binding. We filtered gRNA readouts to only include predefined target regions that resulted in a decrease in a candidate gene’s expression (regression coefficient “beta” column < 0) and excluded loci that were flagged in the “outlier_gene” column. We followed the methodology provided in the paper [25] to aggregate gRNA readouts to gRNA groups targeting the same locus. Target loci with adjusted empirical p-values below 0.05 for any of the measured target genes was labeled activating. Regions that failed to reach that threshold for any genes were labeled neutral. This procedure yielded 432 activating and 5122 neutral regions for Gasperini/K562. The reference nucleotide was the midpoint of a target region.

For Fulco/K562, also a CRISPR-dCas9 dataset, we obtained the published adjusted p-values associated for tested candidate regulatory element-gene pairs (E-G pairs) in K562 [38]. This dataset contains aggregated data from 10 CRISPR-based functional characterization studies [57–66], with the[50, 57–64, 66, 67]vast majority of data points (>99%) generated by perturbation with the CRISPRi-FlowFISH screen, which makes use of CRISPR-dCas9 with an attached KRAB domain, targeting DNase-I hypersensitive sites within 450 kb of 30 selected genes. We used the same procedure as the [38] to exclude any E-G pairs that (i) had less than 80% power to a detect 25% effect on gene expression, or (ii) had a fraction change in gene expression that was positive after CRISPR interference, since it suggests repression. We used a p-value threshold of 0.05 to assign the activating and neutral labels E-G pairs. Candidate regulatory elements that were in at least one E-G pair were assigned to the activating label, while all other elements were assigned the neutral label. This yielded 69 activating and 747 neutral regions for Fulco/K562. The reference nucleotide was the center nucleotide of the element.

All training, testing and analysis was done in the hg19 human genome assembly. Genomic coordinates not in hg19 were converted to hg19 using the liftOver utility from the UCSC genome browser [68], specifically from hg18 for Kheradpour HepG2/K562 datasets and from hg38 for White A549/HepG2/K562 datasets. For all datasets, loci not in chromosomes 1 through 22 or chromosome X were filtered out.

### Feature extraction and preprocessing

ChromActivity uses three classes of features in the models: chromatin mark signals, chromatin mark peak calls and ChromHMM chromatin states. For the chromatin signal and peak call features, we used imputed signal tracks and narrow peak calls on imputed signal tracks, respectively, for the following 12 chromatin marks: DNase, H2A.Z, H3K27ac, H3K27me3, H3K36me3, H3K4me1, H3K4me2, H3K4me3, H3K79me2, H3K9ac, H3K9me3, H4K20me1 [9]. For the chromatin state features, we used chromatin state annotations based on the 25-state ChromHMM model based on imputed data [13, 39]. All of these features were available across 127 cell types.

For each region considered, ChromActivity extracts the signal features within a 2 kb window at 25 base intervals centered around a reference nucleotide associated with the region, yielding 81 features per mark. In our application here, this resulted in 972 intermediate signal features for the 12 marks. For each mark, ChromActivity then applies principal component analysis (PCA) to its 81 signal features and selects the top three principal components. In our application here, the first three principal components explained on average 97% of the variance across training regions. ChromActivity retains the original signal value at the reference nucleotide thus reducing the number of signal features from 81 to 4 per mark. In evaluations and analyses that involved dividing the dataset into training and test partitions, PC component weights were learned from training partitions in each dataset and then applied to the test partitions.

For each chromatin mark peak, ChromActivity includes a binary indicator variable for the presence of the peak at the reference nucleotide. It also includes features corresponding to a one-hot encoding of the 25-chromatin state annotation at the reference nucleotides. Altogether, this procedure yields 85 features used for classification: 36 PCA signal features, 12 original signal value features, 12 chromatin mark peak features and 25 one-hot encoded chromatin state annotations features. All features are standardized (based on the training partition for evaluations involving train and test sets) to have mean zero and a variance of one before training.

### Training, evaluation, and genomewide prediction track generation of the expert models

ChromActivity uses a bagging ensemble of logistic regression classifiers to generate the individual experts, which has the advantages of being robust and providing well-calibrated probability estimates that reflect the class membership of the training data. For each functional characterization dataset, ChromActivity trained an ensemble of classifiers based on the extracted labels and features as described above. Each ensemble contained 100 binary logistic regression classifiers with a L2-norm penalty trained on a random drawing of training data points. The data points were drawn with replacement to obtain the same number of data points as the initial training set, i.e. a bagging ensemble. The regularization strength *C* of the logistic regression classifiers was set to the default value of 1 and assigned label weights of w(y=*activating*) = n_neutral_ and w(y=*neutral*) = 3n_activating_ to the label classes, where *w*(y=*y’*) indicates label class weight for *y’*, and *n_y’_* indicates number of data points of label *y’*. The label weights correspond to an effective label ratio of 0.25 (activating/neutral) across different datasets (Supplementary Figure S3A) instead of a balanced ratio so the resulting score better highlights genomic regions with high regulatory activity potential.

To evaluate the predictive performance of ChromActivity’s expert models on the functional characterization datasets they were trained on, we randomly generated 20 train/test partitions per dataset with a 4:1 train:test ratio, stratified by label to ensure consistent label ratios across the partitions. The models were trained as described above and applied to each test set to obtain AUROC metrics in Supplementary Figure S4.

To produce the genomewide expert score tracks, ChromActivity applied the experts (trained on the entire training data) at 25 bp intervals across the genome to predict the activating class label probability at the center nucleotide in the interval (i.e. 13th nucleotide). ChromActivity produced ChromScore by taking the mean value of the individual expert predictions for each 25 bp interval.

We computed a normalized version of the expert scores for analyses in which the distributions of the expert model scores are directly compared on the same sets of genomic loci (Figure 2B, 2C, Supplementary Figures S5-7). The normalization procedure we implemented was based on quantile normalization. Specifically, to establish the reference distribution, we first computed expert model scores for 10 million randomly selected genomic locations, removing regions in the ENCODE excluded list [56]. We sorted the expert scores and computed the median expert score for each ranked entry. We then computed 1,000 quantile bins of each expert score distribution and generated mappings from the quantile bins to the corresponding median expert scores. Score values from experts are mapped to normalized score values using these mappings. We computed the mean normalized expert score values over all experts to generate the normalized ChromScore track used in Supplementary Figure S7.

### ChromScoreHMM annotations

To generate the ChromScoreHMM annotations, ChromActivity first converts the continuous score tracks associated with expert models into binarized input for ChromHMM (version 1.23). These annotations are generated at 25 bp resolution, corresponding to the resolution of the predictions, instead of the default ChromHMM resolution of 200bp. For the main analysis, the binarization thresholds per score track were set such that the 25 bp bins within the top 2% of model scores were assigned to 1 and the rest were assigned to 0.

We used ChromHMM’s LearnModel subcommand with the following command line flags: -b 25 - n 128 -p 4 -d -1 -lowmem. This configuration corresponds to a score bin size of 25 bases, using 128 randomly selected cell type and chromosome combinations per Baum-Welch training iteration, 4 threads running the standard Baum-Welch algorithm, with the change in estimated log-likelihood stopping criterion disabled and reduced memory usage mode. The number of chromatin states for the main analysis was set to 15. Emission and transmission parameters of the model are shown in Supplementary Figure S9.

To determine the number of states and the binarization threshold we ran models with the number of chromatin states set to 10, 15, 25 and binarization thresholds of top 1%, 2%, 5% and 10%. We focused on a 15-state model as it provided a good balance between model expressivity and interpretability for multiple values of the binarization threshold. The binarization threshold presented a tradeoff: a higher binarization threshold risks missing a larger number of true regulatory sites or evidence that a regulatory site is supported by multiple expert’s top predictions, while a lower binarization threshold could overassign the genome into regulatory states (Supplementary Figure S20). We opted to use a binarization threshold of 2%, which provided a reasonable tradeoff with approximately 9.3% of the genome in a cell type on average in ChromScoreHMM states associated with at least one expert (Figure 3) and 5.0% of the 25 bp intervals in the genome were above the binarization threshold in two or more experts (Supplementary Figure S20).

### ChromScoreHMM overlap fold enrichments

We computed overlap fold enrichments using ChromHMM’s OverlapEnrichment command with command line flag -b 25. CpG island coordinates and RefSeq gene coordinates [41] were the ones included with ChromHMM (version 1.23), originally downloaded from the UCSC genome browser. RefSeq annotations were the version available on July 26, 2015. RepeatMasker [45] repeat element coordinates and PhastCons 100-way conserved element annotations [44] were obtained from the UCSC Genome Browser [40, 69]. CTCF motif instances were obtained from HOMER known motifs (version 191020) [42]. GERP++ conserved element annotations were obtained from http://mendel.stanford.edu/SidowLab/downloads/gerp [43].

### Analysis of expert score and ChromScore distributions

#### Pairwise expert score correlations

We computed pairwise Pearson correlations between pairs of expert scores at 500,000 randomly selected bases of the genome, excluding any region on the ENCODE excluded regions list v2 for hg19 [56].

#### Chromatin state score distributions

To determine chromatin state score distributions for the 25-state ChromHMM annotations [13, 39], we sampled 2.5 million loci from the genome excluding those in ENCODE excluded regions v2 as above and extracted their chromatin states and associated scores.

#### Cluster heatmap of ChromScores across cell types and tissue group correlations

To generate a cluster heatmap of scores across cell types, we randomly selected 20,000 bases from the genome among those for which ChromScore showed a difference of at least 0.25 between at least one of the 127 cell types. We filtered for score differences to highlight genomic loci with different regulatory activity potential across cell types. Roadmap Epigenomics tissue groupings were obtained from the metadata section of the Roadmap Epigenomics data portal [9]. Loci were clustered using the euclidean average linkage metric implemented in scipy.cluster.hierarchy.linkage in the SciPy package [70]. We excluded the “ENCODE2012” and “Other” tissue groupings when computing the mean ChromScore correlations within and across tissue groups.

### Evaluating ChromScore cell type generalization performance

To estimate the generalization performance of ChromScore in unseen cell types, we trained five modified versions of the model, one for each cell type with characterization data available. Each version was constructed such that it did not have access to training data in one particular cell type (i.e. one of A549, GM12878, HeLaS3, HepG2 or K562). To evaluate predictive performance for a dataset of a particular cell type, we used the version of the model with that cell type removed. In addition to holding out cell types, we also spatially partitioned the genome into 5 kb chunks and assigned each chunk to the training or testing partition with probability 0.25 (i.e. 3:1 train:test ratio). We repeated this process 20 times per dataset.

We compared the performance of ChromScore to a set of baselines and existing scores. The baselines included the individual imputed chromatin mark signals (DNase, H2A.Z, H3K27ac, H3K27me3, H3K36me3, H3K4me1, H3K4me2, H3K4me3, H3K79me2, H3K9ac, H3K9me3, and H4K20me1) in the matched cell types obtained from the Roadmap Epigenomics compendium. In addition, they included a simple chromatin state baseline model, which generated a single score track for each cell type by mapping a chromatin state annotation at a specific position to the average fraction of positive labels within the training partition for each dataset.

We also compared ChromScore to various cell type-specific and non-cell type-specific external scores that integrate different epigenomic datasets and in some cases with other annotations, specifically FunLDA [47], GenoSkyline Plus [48], LINSIGHT [50], CADD [49, 51] and GenoNet/GenoNet-U [27]. Precomputed FunLDA scores were downloaded from http://www.funlda.com/download. GenoSkyline Plus annotations were obtained from http://zhaocenter.org/GenoSkyline. LINSIGHT annotation was downloaded from http://compgen.cshl.edu/LINSIGHT. CADD v1.4 scores were obtained from https://cadd.gs.washington.edu. We used the browser track (hg19) version of the CADD scores, which are based on the highest scoring single nucleotide variant for each genomic position as described in https://github.com/kircherlab/CADD-browserTracks.

GenoNet used two distinct models, a supervised version (“GenoNet”) which was trained on MPRA data [33, 71] and was only applied to the three cell types K562, HepG2, GM12878 and an unsupervised version (“GenoNet-U”) which did not use any functional characterization data and was applied to the remaining 124 Roadmap Epigenome cell types. Precomputed GenoNet scores for K562, HepG2 and GM12878 and precomputed Genonet-U scores for the remaining Roadmap Epigenomics cell type were obtained from https://zenodo.org/record/3336208 [27, 72]. Precomputed GenoNet-U scores in K562, HepG2, GM12878 were not available, and instead we computed using a custom script based on the description of the method. The output of our implementation was confirmed to produce nearly identical predictions (Pearson correlation > 0.99) to the GenoNet-U scores in all 124 of Roadmap epigenome cell types for which it was available.

### Expression analyses around TSSs

We downloaded the RPKM expression matrix for protein coding genes for 56 Roadmap Epigenomes from the Roadmap Epigenomics data portal (https://egg2.wustl.edu/roadmap/data/byDataType/rna/expression/57epigenomes.RPKM.pc.gz), along with the corresponding Ensembl gene annotations (Ensembl v65, hg19) [73]. For each cell type, we categorized the genes into high expression (defined as log_2_(RPKM+1) > 1) and low expression (defined as log_2_(RPKM+1) < 0.01) genes. Across the 56 cell types, 62% of genes were categorized as high expression and 15% of genes were categorized as low expression on average.

To investigate expression of nearby genes for ChromScoreHMM states, we identified the ChromScoreHMM states within 24 kb windows centered around the TSSs of the genes, sampled at 200 base intervals. For genes on the negative strand, we flipped the position indices so that positive offset values always corresponded to the direction of the gene body. We computed log_2_(RPKM+1) values for each gene based on the Roadmap Epigenomics RPKM expression matrix for protein coding genes. For each ChromScoreHMM state and position offset, we then computed the mean log_2_(RPKM+1) value across genes and cell types.

For the ChromScore expression correlations analysis, we first computed ChromScore and individual expert model scores within a 24 kb window centered around TSSs in all cell types with expression data available. We also extracted chromatin mark signal values for the same windows for comparison. We mirrored the score windows for the genes on the negative strand around the TSS to align the upstream segments and the gene bodies. We then computed Pearson correlations between scores or signal values and log expression with pseudocount (log_2_(RPKM+1)) for each 25 bp interval centered around the TSS and averaged them over the cell types.

### ChromScore repeat element enrichments

We downloaded RepeatMasker (Smit, Hubley and Green, 2013) repeat elements from the UCSC Genome Browser (Karolchik et al., 2004; Navarro Gonzalez et al., 2021), using the repClass column to identify the LINE, SINE and LTR elements and the repFamily column to identify the ERV elements. We randomly selected 1 million nucleotides from hg19 on chromosomes 1 through 22 and chromosome X, determined if they overlapped a repeat element with the bedtools intersect command, and computed ChromScore and expert prediction scores for each plasmid-based and CRISPR-based expert. Mean plasmid and CRISPR scores were obtained by taking the mean score of the respective experts at each nucleotide. We grouped the scores within 200 quantiles (i.e. each quantile representing 0.5% of the nucleotides) and computed fold enrichments for each quantile for each repeat type compared to the genome background.

## Declarations

### Ethics approval and consent to participate

Not applicable.

### Consent for publication

Not applicable.

### Availability of data and materials

The ChromActivity software and links to the ChromScoreHMM and ChromScore annotations (hg19 and hg38 liftOver) for the 127 Roadmap cell types are available on https://github.com/ernstlab/ChromActivity. Signal and score tracks were processed using pyBigWig v0.3.18 [74] and UCSC utilities v369 [68, 75]. Genomic coordinates were mapped across genome assemblies using liftOver v369 [68, 75]. Analyses involving genome intervals used BedTools v2.30.0 [76], pyBedTools v0.9.0 [77], bedops v2.4.41 [78] packages. Hierarchical clustering used in heatmap visualizations is implemented in SciPy [70]. scikit-learn v1.1.2 [79] was used for data preprocessing, model training, inference and evaluations. ChromScoreHMM annotations were generated using ChromHMM v.1.23 [13, 14], downloaded from https://ernstlab.biolchem.ucla.edu/ChromHMM. We used matplotlib [80] and Seaborn [81] for plotting and visualization. All other packages were obtained from the conda-forge and bioconda [82] repositories.

### Authors’ contributions

TD designed and implemented ChromActivity, compiled and curated input datasets, performed analyses and wrote the paper. JE conceived the study, designed ChromActivity, supervised the work, and wrote the paper. All authors read and approved the final manuscript.

## Acknowledgements

We thank Heather Han for conducting some related preliminary work and members of the Ernst lab for insightful discussions. We acknowledge the ENCODE and Roadmap Epigenomics Consortia and the individual labs that generated the data that we used.

## References

1. Chatterjee S, Ahituv N. Gene Regulatory Elements, Major Drivers of Human Disease. Annu Rev Genomics Hum Genet. 2017;18:45–63.

2. Lambert SA, Jolma A, Campitelli LF, Das PK, Yin Y, Albu M, et al. The Human Transcription Factors. Cell. 2018;172:650–65.

3. Gasperini M, Tome JM, Shendure J. Towards a comprehensive catalogue of validated and target-linked human enhancers. Nat Rev Genet. 2020;21:292–310.

4. Ernst J, Kheradpour P, Mikkelsen TS, Shoresh N, Ward LD, Epstein CB, et al. Mapping and analysis of chromatin state dynamics in nine human cell types. Nature. 2011;473:43–9.

5. Maurano MT, Humbert R, Rynes E, Thurman RE, Haugen E, Wang H, et al. Systematic Localization of Common Disease-Associated Variation in Regulatory DNA. Science. 2012;337:1190–5.

6. Barski A, Cuddapah S, Cui K, Roh T-Y, Schones DE, Wang Z, et al. High-resolution profiling of histone methylations in the human genome. Cell. 2007;129:823–37.

7. The ENCODE Project Consortium. An integrated encyclopedia of DNA elements in the human genome. Nature. 2012;489:57–74.

8. Thurman RE, Rynes E, Humbert R, Vierstra J, Maurano MT, Haugen E, et al. The accessible chromatin landscape of the human genome. Nature. 2012;489:75–82.

9. Roadmap Epigenomics Consortium, Kundaje A, Meuleman W, Ernst J, Bilenky M, Yen A, et al. Integrative analysis of 111 reference human epigenomes. Nature. 2015;518:317–30.

10. Stunnenberg HG, Abrignani S, Adams D, de Almeida M, Altucci L, Amin V, et al. The International Human Epigenome Consortium: A Blueprint for Scientific Collaboration and Discovery. Cell. 2016;167:1145–9.

11. Buenrostro JD, Giresi PG, Zaba LC, Chang HY, Greenleaf WJ. Transposition of native chromatin for fast and sensitive epigenomic profiling of open chromatin, DNA-binding proteins and nucleosome position. Nat Methods. 2013;10:1213–8.

12. Zhang Y, Liu T, Meyer CA, Eeckhoute J, Johnson DS, Bernstein BE, et al. Model-based Analysis of ChIP-Seq (MACS). Genome Biol. 2008;9:R137.

13. Ernst J, Kellis M. ChromHMM: automating chromatin-state discovery and characterization. Nat Methods. 2012;9:215–6.

14. Ernst J, Kellis M. Chromatin-state discovery and genome annotation with ChromHMM. Nat Protoc. 2017;12:2478–92.

15. Hoffman MM, Buske OJ, Wang J, Weng Z, Bilmes JA, Noble WS. Unsupervised pattern discovery in human chromatin structure through genomic segmentation. Nat Methods. 2012;9:473–6.

16. Libbrecht MW, Chan RCW, Hoffman MM. Segmentation and genome annotation algorithms for identifying chromatin state and other genomic patterns. PLOS Comput Biol. 2021;17:e1009423.

17. Andersson R, Gebhard C, Miguel-Escalada I, Hoof I, Bornholdt J, Boyd M, et al. An atlas of active enhancers across human cell types and tissues. Nature. 2014;507:455–61.

18. Andersson R, Sandelin A. Determinants of enhancer and promoter activities of regulatory elements. Nat Rev Genet. 2020;21:71–87.

19. Klein JC, Agarwal V, Inoue F, Keith A, Martin B, Kircher M, et al. A systematic evaluation of the design and context dependencies of massively parallel reporter assays. Nat Methods. 2020;17:1083–91.

20. Chen PB, Fiaux PC, Zhang K, Li B, Kubo N, Jiang S, et al. Systematic discovery and functional dissection of enhancers needed for cancer cell fitness and proliferation. Cell Rep. 2022;41:111630.

21. Inoue F, Ahituv N. Decoding enhancers using massively parallel reporter assays. Genomics. 2015;106:159–64.

22. Melnikov A, Murugan A, Zhang X, Tesileanu T, Wang L, Rogov P, et al. Systematic dissection and optimization of inducible enhancers in human cells using a massively parallel reporter assay. Nat Biotechnol. 2012;30:271–7.

23. Gallego Romero I, Lea AJ. Leveraging massively parallel reporter assays for evolutionary questions. Genome Biol. 2023;24:26.

24. Arnold CD, Gerlach D, Stelzer C, Boryń ŁM, Rath M, Stark A. Genome-wide quantitative enhancer activity maps identified by STARR-seq. Science. 2013;339:1074–7.

25. Gasperini M, Hill AJ, McFaline-Figueroa JL, Martin B, Kim S, Zhang MD, et al. A Genome wide Framework for Mapping Gene Regulation via Cellular Genetic Screens. Cell. 2019;176:377–390.e19.

26. Chong ZX, Yeap SK, Ho WY. Transfection types, methods and strategies: a technical review. PeerJ. 2021;9:e11165.

27. He Z, Liu L, Wang K, Ionita-Laza I. A semi-supervised approach for predicting cell-type specific functional consequences of non-coding variation using MPRAs. Nat Commun. 2018;9:5199.

28. Sethi A, Gu M, Gumusgoz E, Chan L, Yan K-K, Rozowsky J, et al. Supervised enhancer prediction with epigenetic pattern recognition and targeted validation. Nat Methods. 2020;17:807–14.

29. Kreimer A, Zeng H, Edwards MD, Guo Y, Tian K, Shin S, et al. Predicting gene expression in massively parallel reporter assays: A comparative study. Hum Mutat. 2017;38:1240–50.

30. Kreimer A, Yan Z, Ahituv N, Yosef N. Meta-analysis of massively parallel reporter assays enables prediction of regulatory function across cell types. Hum Mutat. 2019;40:1299–313.

31. Zhou J, Theesfeld CL, Yao K, Chen KM, Wong AK, Troyanskaya OG. Deep learning sequence-based ab initio prediction of variant effects on expression and disease risk. Nat Genet. 2018;50:1171–9.

32. Movva R, Greenside P, Marinov GK, Nair S, Shrikumar A, Kundaje A. Deciphering regulatory DNA sequences and noncoding genetic variants using neural network models of massively parallel reporter assays. PloS One. 2019;14:e0218073.

33. Kheradpour P, Ernst J, Melnikov A, Rogov P, Wang L, Zhang X, et al. Systematic dissection of regulatory motifs in 2000 predicted human enhancers using a massively parallel reporter assay. Genome Res. 2013;23:800–11.

34. Ernst J, Melnikov A, Zhang X, Wang L, Rogov P, Mikkelsen TS, et al. Genome-scale high resolution mapping of activating and repressive nucleotides in regulatory regions. Nat Biotechnol. 2016;34:1180–90.

35. Muerdter F, Boryń ŁM, Woodfin AR, Neumayr C, Rath M, Zabidi MA, et al. Resolving systematic errors in widely used enhancer activity assays in human cells. Nat Methods. 2018;15:141–9.

36. Wang X, He L, Goggin SM, Saadat A, Wang L, Sinnott-Armstrong N, et al. High-resolution genome-wide functional dissection of transcriptional regulatory regions and nucleotides in human. Nat Commun. 2018;9:5380.

37. Lee D, Shi M, Moran J, Wall M, Zhang J, Liu J, et al. STARRPeaker: uniform processing and accurate identification of STARR-seq active regions. Genome Biol. 2020;21:298.

38. Fulco CP, Nasser J, Jones TR, Munson G, Bergman DT, Subramanian V, et al. Activity-by-contact model of enhancer–promoter regulation from thousands of CRISPR perturbations. Nat Genet. 2019;51:1664–9.

39. Ernst J, Kellis M. Large-scale imputation of epigenomic datasets for systematic annotation of diverse human tissues. Nat Biotechnol. 2015;33:364–76.

40. Karolchik D, Hinrichs AS, Furey TS, Roskin KM, Sugnet CW, Haussler D, et al. The UCSC Table Browser data retrieval tool. Nucleic Acids Res. 2004;32 Database issue:D493–496.

41. O’Leary NA, Wright MW, Brister JR, Ciufo S, Haddad D, McVeigh R, et al. Reference sequence (RefSeq) database at NCBI: current status, taxonomic expansion, and functional annotation. Nucleic Acids Res. 2016;44:D733–745.

42. Heinz S, Benner C, Spann N, Bertolino E, Lin YC, Laslo P, et al. Simple combinations of lineage-determining transcription factors prime cis-regulatory elements required for macrophage and B cell identities. Mol Cell. 2010;38:576–89.

43. Davydov EV, Goode DL, Sirota M, Cooper GM, Sidow A, Batzoglou S. Identifying a High Fraction of the Human Genome to be under Selective Constraint Using GERP++. PLOS Comput Biol. 2010;6:e1001025.

44. Siepel A, Bejerano G, Pedersen JS, Hinrichs AS, Hou M, Rosenbloom K, et al. Evolutionarily conserved elements in vertebrate, insect, worm, and yeast genomes. Genome Res. 2005;15:1034–50.

45. Smit A, Hubley R, Green P. RepeatMasker Open-4.0. 2013.

46. Vu H, Ernst J. Universal annotation of the human genome through integration of over a thousand epigenomic datasets. Genome Biol. 2022;23:9.

47. Backenroth D, He Z, Kiryluk K, Boeva V, Petukhova L, Khurana E, et al. FUN-LDA: A Latent Dirichlet Allocation Model for Predicting Tissue-Specific Functional Effects of Noncoding Variation: Methods and Applications. Am J Hum Genet. 2018;102:920–42.

48. Lu Q, Powles RL, Abdallah S, Ou D, Wang Q, Hu Y, et al. Systematic tissue-specific functional annotation of the human genome highlights immune-related DNA elements for late onset Alzheimer’s disease. PLOS Genet. 2017;13:e1006933.

49. Rentzsch P, Witten D, Cooper GM, Shendure J, Kircher M. CADD: predicting the deleteriousness of variants throughout the human genome. Nucleic Acids Res. 2019;47:D886– 94.

50. Huang Y-F, Gulko B, Siepel A. Fast, scalable prediction of deleterious noncoding variants from functional and population genomic data. Nat Genet. 2017;49:618–24.

51. Kircher M, Witten DM, Jain P, O’Roak BJ, Cooper GM, Shendure J. A general framework for estimating the relative pathogenicity of human genetic variants. Nat Genet. 2014;46:310–5.

52. Bannert N, Kurth R. Retroelements and the human genome: New perspectives on an old relation. Proc Natl Acad Sci. 2004;101 suppl_2:14572–9.

53. Criscione SW, Zhang Y, Thompson W, Sedivy JM, Neretti N. Transcriptional landscape of repetitive elements in normal and cancer human cells. BMC Genomics. 2014;15:583.

54. Ali A, Han K, Liang P. Role of Transposable Elements in Gene Regulation in the Human Genome. Life. 2021;11:118.

55. Zhang X-O, Pratt H, Weng Z. Investigating the Potential Roles of SINEs in the Human Genome. Annu Rev Genomics Hum Genet. 2021;22:199–218.

56. Amemiya HM, Kundaje A, Boyle AP. The ENCODE Blacklist: Identification of Problematic Regions of the Genome. Sci Rep. 2019;9:9354.

57. Thakore PI, D’Ippolito AM, Song L, Safi A, Shivakumar NK, Kabadi AM, et al. Highly specific epigenome editing by CRISPR-Cas9 repressors for silencing of distal regulatory elements. Nat Methods. 2015;12:1143–9.

58. Xu J, Shao Z, Li D, Xie H, Kim W, Huang J, et al. Developmental control of polycomb subunit composition by GATA factors mediates a switch to non-canonical functions. Mol Cell. 2015;57:304–16.

59. Fulco CP, Munschauer M, Anyoha R, Munson G, Grossman SR, Perez EM, et al. Systematic mapping of functional enhancer–promoter connections with CRISPR interference. Science. 2016;354:769–73.

60. Ulirsch JC, Nandakumar SK, Wang L, Giani FC, Zhang X, Rogov P, et al. Systematic Functional Dissection of Common Genetic Variation Affecting Red Blood Cell Traits. Cell. 2016;165:1530–45.

61. Wakabayashi A, Ulirsch JC, Ludwig LS, Fiorini C, Yasuda M, Choudhuri A, et al. Insight into GATA1 transcriptional activity through interrogation of cis elements disrupted in human erythroid disorders. Proc Natl Acad Sci U S A. 2016;113:4434–9.

62. Klann TS, Black JB, Chellappan M, Safi A, Song L, Hilton IB, et al. CRISPR–Cas9 epigenome editing enables high-throughput screening for functional regulatory elements in the human genome. Nat Biotechnol. 2017;35:561–8.

63. Liu SJ, Horlbeck MA, Cho SW, Birk HS, Malatesta M, He D, et al. CRISPRi-based genome scale identification of functional long noncoding RNA loci in human cells. Science. 2017;355:eaah7111.

64. Xie S, Duan J, Li B, Zhou P, Hon GC. Multiplexed Engineering and Analysis of Combinatorial Enhancer Activity in Single Cells. Mol Cell. 2017;66:285–299.e5.

65. Huang J, Li K, Cai W, Liu X, Zhang Y, Orkin SH, et al. Dissecting super-enhancer hierarchy based on chromatin interactions. Nat Commun. 2018;9:943.

66. Qi Z, Xie S, Chen R, Aisa HA, Hon GC, Guan Y. Tissue-specific Gene Expression Prediction Associates Vitiligo with SUOX through an Active Enhancer. 2018;:337196.

67. Engreitz JM, Haines JE, Perez EM, Munson G, Chen J, Kane M, et al. Local regulation of gene expression by lncRNA promoters, transcription and splicing. Nature. 2016;539:452–5.

68. Kent WJ, Zweig AS, Barber G, Hinrichs AS, Karolchik D. BigWig and BigBed: enabling browsing of large distributed datasets. Bioinformatics. 2010;26:2204–7.

69. Navarro Gonzalez J, Zweig AS, Speir ML, Schmelter D, Rosenbloom KR, Raney BJ, et al. The UCSC Genome Browser database: 2021 update. Nucleic Acids Res. 2021;49:D1046–57.

70. Virtanen P, Gommers R, Oliphant TE, Haberland M, Reddy T, Cournapeau D, et al. SciPy 1.0: fundamental algorithms for scientific computing in Python. Nat Methods. 2020;17:261–72.

71. Tewhey R, Kotliar D, Park DS, Liu B, Winnicki S, Reilly SK, et al. Direct Identification of Hundreds of Expression-Modulating Variants using a Multiplexed Reporter Assay. Cell. 2016;165:1519–29.

72. Ionita-Laza I. GenoNet scores for human genome assembly GRCh37. 2019.

73. Cunningham F, Allen JE, Allen J, Alvarez-Jarreta J, Amode MR, Armean IM, et al. Ensembl 2022. Nucleic Acids Res. 2022;50:D988–95.

74. Ryan D, Gökçen Eraslan, Grüning B, Betts E, Ramirez F, Nezar Abdennur, et al. deeptools/pyBigWig: 0.3.18. 2021.

75. Kent WJ, Sugnet CW, Furey TS, Roskin KM, Pringle TH, Zahler AM, et al. The human genome browser at UCSC. Genome Res. 2002;12:996–1006.

76. Quinlan AR, Hall IM. BEDTools: a flexible suite of utilities for comparing genomic features. Bioinformatics. 2010;26:841–2.

77. Dale RK, Pedersen BS, Quinlan AR. Pybedtools: a flexible Python library for manipulating genomic datasets and annotations. Bioinforma Oxf Engl. 2011;27:3423–4.

78. Neph S, Kuehn MS, Reynolds AP, Haugen E, Thurman RE, Johnson AK, et al. BEDOPS: high-performance genomic feature operations. Bioinforma Oxf Engl. 2012;28:1919–20.

79. Pedregosa F, Varoquaux G, Gramfort A, Michel V, Thirion B, Grisel O, et al. Scikit-learn: Machine Learning in Python. J Mach Learn Res. 2011;12 Oct:2825–30.

80. Hunter JD. Matplotlib: A 2D Graphics Environment. Comput Sci Eng. 2007;9:90–5.

81. Waskom M. seaborn: statistical data visualization. J Open Source Softw. 2021;6:3021.

82. Grüning B, Dale R, Sjödin A, Chapman BA, Rowe J, Tomkins-Tinch CH, et al. Bioconda: sustainable and comprehensive software distribution for the life sciences. Nat Methods. 2018;15:475–6.

