## Supplementary Figures S1-S20 and Supplementary Table 1 for "Integrative epigenomic and functional characterization assay based annotation of regulatory activity across diverse human cell types"

Supplementary Figure S1: Detailed schematic of the ChromActivity framework.

Supplementary Figure S2: Color legends for ChromScoreHMM states and the ChromHMM imputed 25-state model.

Supplementary Figure S3: Genomic coverage of functional characterization datasets.

Supplementary Figure S4: Predictive performance of ChromScore expert models in held-out loci of the same functional characterization dataset.

Supplementary Figure S5: Mean normalized expert scores per chromatin state.

Supplementary Figure S6: Boxplots of mean normalized expert scores for each chromatin state, distributions across cell types.

Supplementary Figure S7: ChromScoreHMM emission parameters and mean model scores for the 15-state model.

Supplementary Figure S8: ChromScoreHMM emission parameters and chromatin mark peak enrichments for the 15-state model.

Supplementary Figure S9: ChromScoreHMM emission and transition parameters for the 15-state model.

Supplementary Figure S10: ChromScoreHMM state proportions around transcription start sites.

Supplementary Figure S11: ChromScoreHMM gene expression patterns around transcription start sites.

Supplementary Figure S12: ChromScoreHMM overlap enrichments around TSSs of high expression and low expression genes.

Supplementary Figure S13: Predictive performance of ChromScore, external models and baselines.

Supplementary Figure S14: Predictive performance of ChromScore and individual experts.

Supplementary Figure S15: Predictive performance of ChromScore, external models and baselines in CRISPR and plasmid-based functional characterization datasets.

Supplementary Figure S16: Spearman correlations between ChromScore and chromatin mark signal tracks.

Supplementary Figure S17: Chromatin state fold enrichments of top ChromScore and DNase-I hypersensitive regions.

Supplementary Figure S18: Repeat element fold enrichments for ChromScore, mean plasmid-based expert score, mean CRISPR-based expert score and DNase signal quantiles.

Supplementary Figure S19: Cell type averages of Pearson correlations between gene expression and ChromScore, individual chromatin mark signals, expert scores.

Supplementary Figure S20: Percentage of genome above the binarization threshold by one or more experts.

### Supplementary Tables

Supplementary Table 1: Overview of the functional characterization datasets.

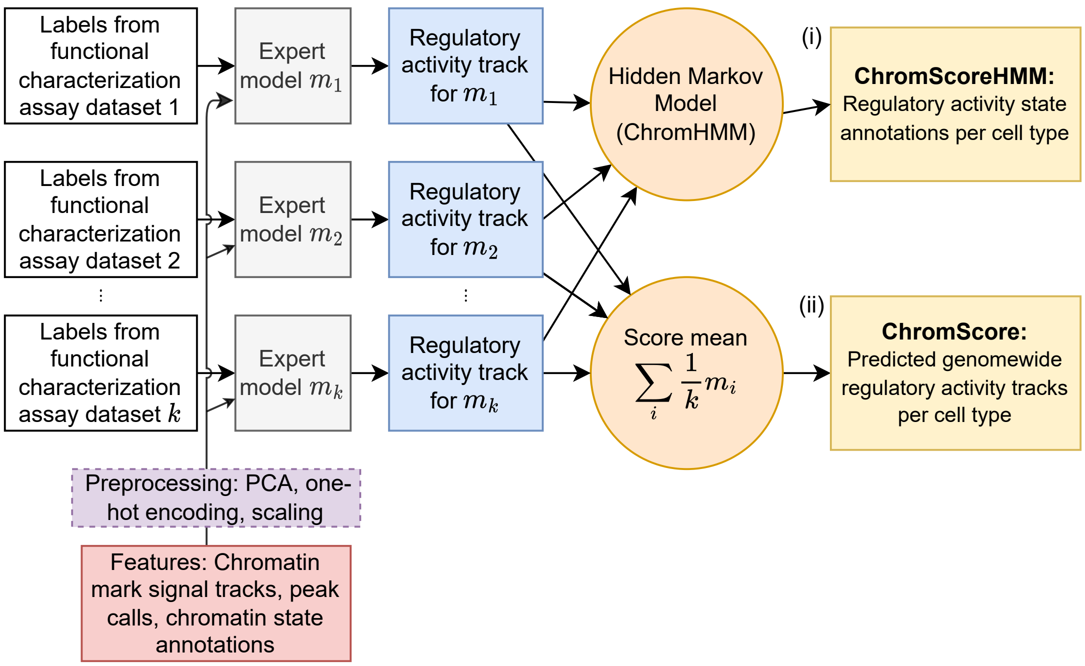


**Supplementary Figure S1: Detailed schematic of the ChromActivity framework.** As summarized in Figure 1A, ChromActivity takes as input regulatory activity labels (white blocks, upper left) for tested genomic regions from multiple functional characterization datasets. ChromActivity also takes as input features based on chromatin mark signals, peak calls and chromatin state assignments for the input regions (red block, lower left) which it preprocesses (purple block, lower left) and uses along with the labels to train a separate classifier ("expert") for each functional characterization dataset ("expert models", gray blocks). Each expert model provides a predicted genomewide regulatory activity score track specific to a functional characterization dataset (blue blocks). ChromActivity then uses these score tracks to generate two complementary outputs (yellow blocks, right): **(i)** ChromScoreHMM annotations, which are annotations of the genome into states based on combinatorial and spatial patterns in the predicted regulatory activity tracks and **(ii)** ChromScore, which is a composite regulatory activity score track based on the mean of individual expert model score tracks.

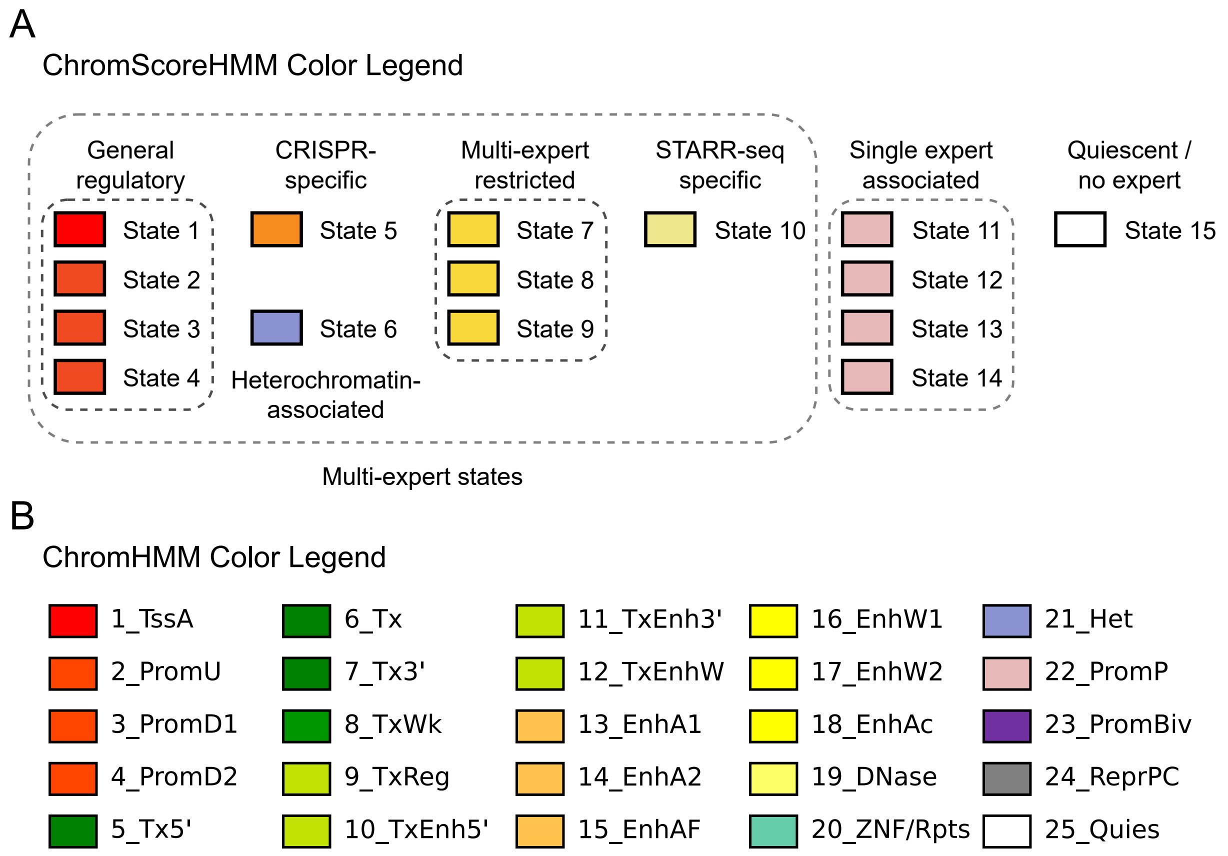


**Supplementary Figure S2: Color legends for ChromScoreHMM states and the ChromHMM imputed 25-state model. (A)** The color legend shows the ChromScoreHMM states grouped into seven color groups for visualization purposes. The groups and their associated states are as follows: General regulatory (States 1, 2, 3 and 4), which are broadly associated with regulatory activity across the majority of experts; CRISPR-specific (State 5), which is associated with CRISPR-based experts; Heterochromatin-associated (State 6), which is highly enriched for a heterochromatin chromatin state; Multi-expert restricted (States 7, 8 and 9), which are associated with two or three experts; STARR-seq specific (State 10), which is associated with a subset of STARR-seq-based experts; Single expert associated (States 11, 12, 13 and 14), which are emission parameters dominated by a single expert; Quiescent/no expert (State 15), which is associated with minimal predicted regulatory activity in all experts. **(B)** Color legend for the ChromHMM imputed 25-state model [1, 2].

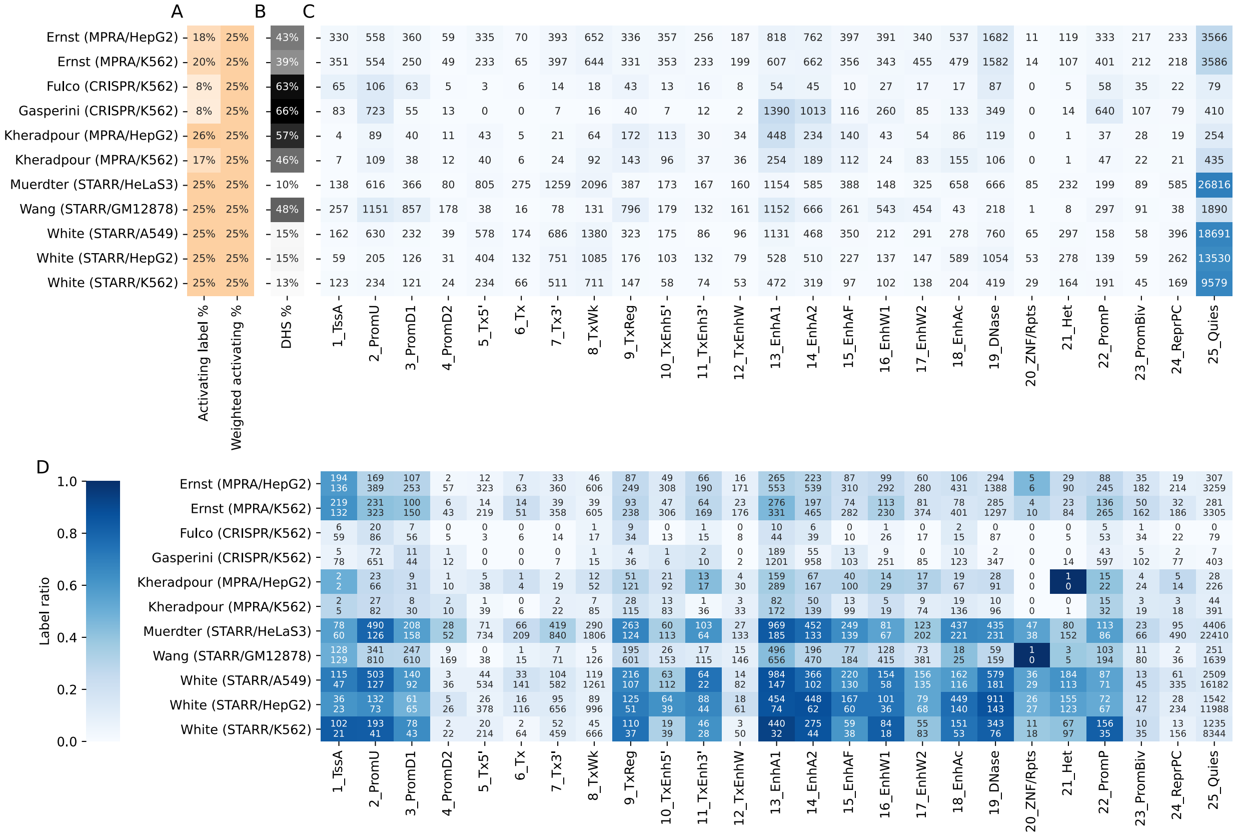


**Supplementary Figure S3: Genomic coverage of functional characterization datasets. (A)** Activating label %: Fraction of dataset loci labeled "activating". Weighted activating %: Effective activating label fraction after label class weighting (Methods). Weighting was not necessary for STARR-seq datasets as neutral loci were selected with a 3-to-1 neutral to activating label ratio (Methods). **(B)** DHS %: Fraction of dataset loci overlapping peaks of DNase in corresponding cell type. Peak calls were based on imputed data obtained from Roadmap Epigenomics [3]. **(C)** Total number of dataset loci in each chromatin state. **(D)** Number of dataset loci labeled "activating" (top) or "neutral" (bottom) in each chromatin state. Heatmap color indicates label ratio as indicated by the color legend on the left.

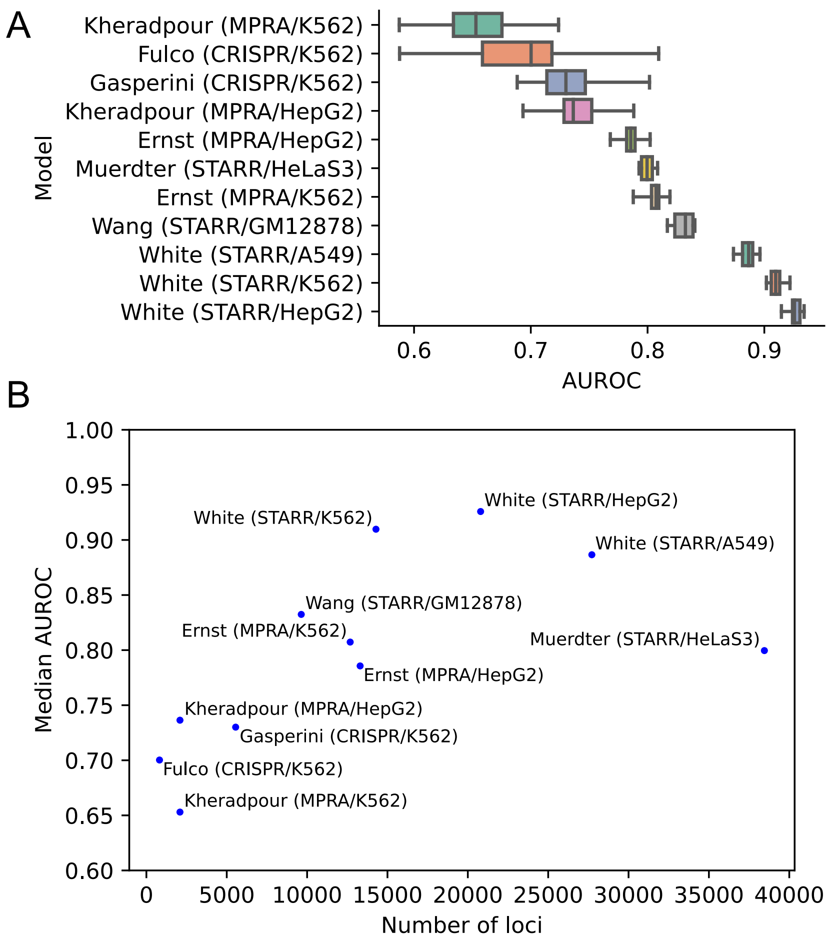


**Supplementary Figure S4: Predictive performance of ChromScore expert models in held-out loci of the same functional characterization dataset. (A)** Area under receiver operator characteristic curve (AUROC) distribution across 20 random permutation cross-validations for each expert model at predicting held-out loci in the same functional characterization dataset (Methods). The box represents quartiles and whiskers indicate maximum and minimum AUROCs across the train/test shuffles. **(B)** Median AUROCs for each expert across the train/test shuffles vs. number of genomic loci used in training (Spearman correlation 0.75).

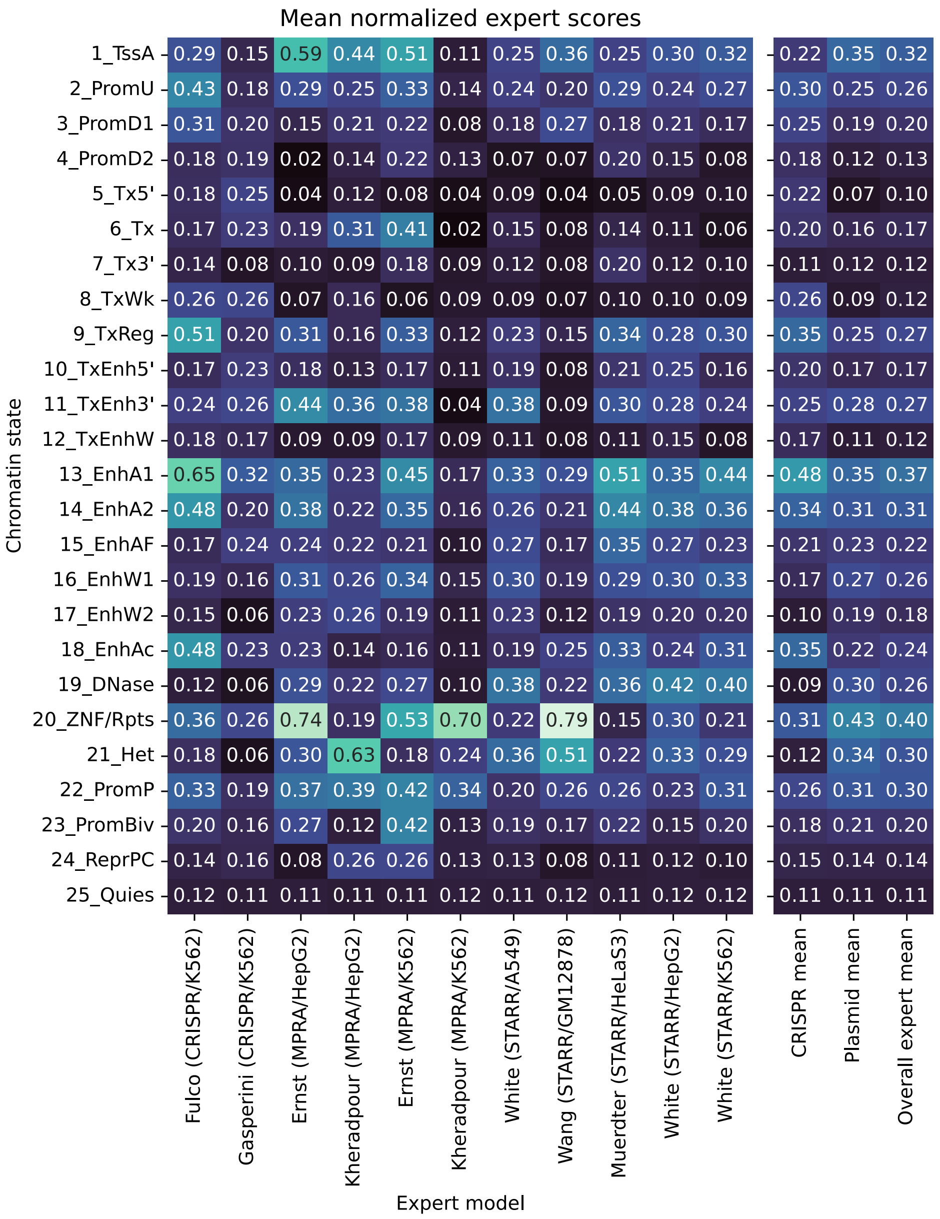


**Supplementary Figure S5: Mean normalized expert scores per chromatin state.** Rows correspond to chromatin states in the 25-state ChromHMM model based on imputed marks [1, 2, 4]. First 11 columns correspond to individual expert models. Values correspond to the mean normalized expert score averaged over the 127 cell types. The rightmost three columns correspond to the mean of the normalized scores across CRISPR-based experts, plasmid-based experts and all experts. See Methods for a description of the normalization procedure.

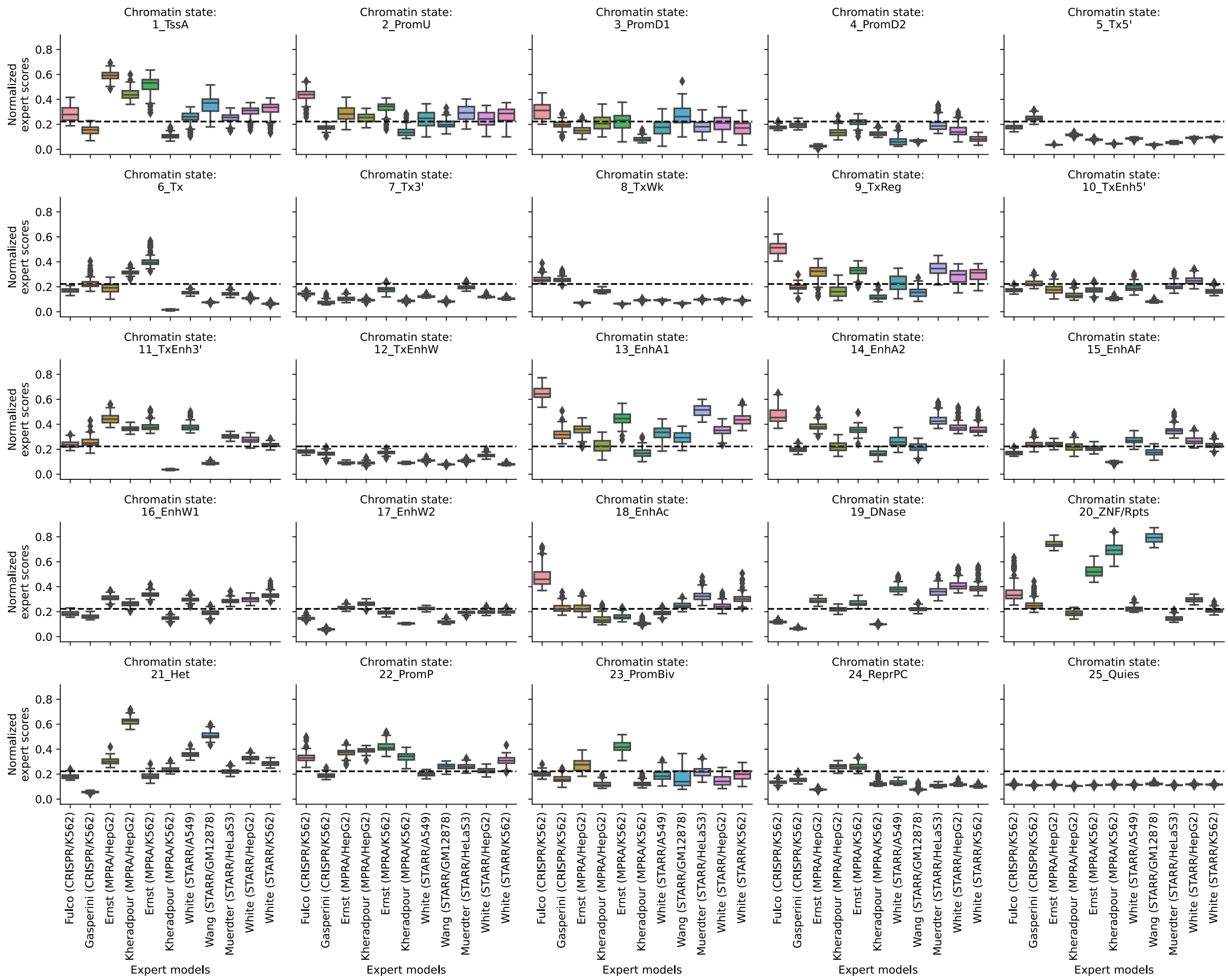


**Supplementary Figure S6: Boxplots of mean normalized expert scores for each chromatin state, distributions across cell types.** Each subplot corresponds to a chromatin state in a 25-state ChromHMM model. Each boxplot within a subplot indicates the distribution of mean normalized expert scores across cell types for the indicated expert and chromatin state (Methods). Horizontal line in the middle of the box indicates median, the box indicates upper and lower quartiles. Dashed line indicates average normalized score across chromatin states.

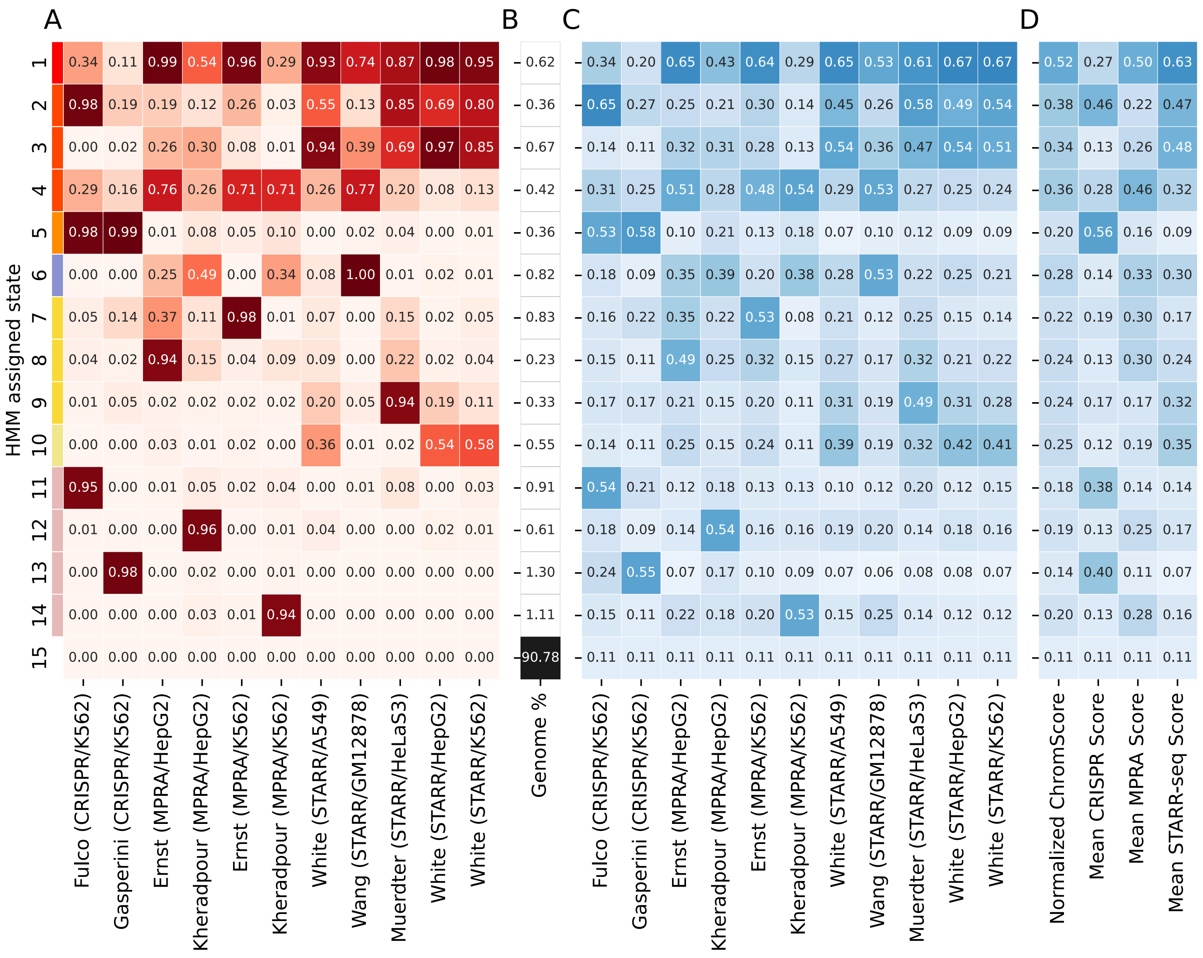


**Supplementary Figure S7: ChromScoreHMM emission parameters and mean model scores for the 15-state model. (A):** Emission parameters of ChromScoreHMM model (see Figure 3 for details). **(B)** Percentage of the genome assigned to each ChromScoreHMM state. **(C)** Mean normalized scores (Methods) for each expert model in each ChromScoreHMM state. **(D)** Mean normalized scores across ChromScoreHMM states (Methods). Normalized ChromScore refers to the mean normalized expert score track (Methods). Mean CRISPR, MPRA and STARR-seq scores refer to scores generated by taking the ensemble averages of normalized score tracks for expert models trained on CRISPR, MPRA and STARR-seq datasets respectively (Methods). Scores and percentages shown are medians across cell types.

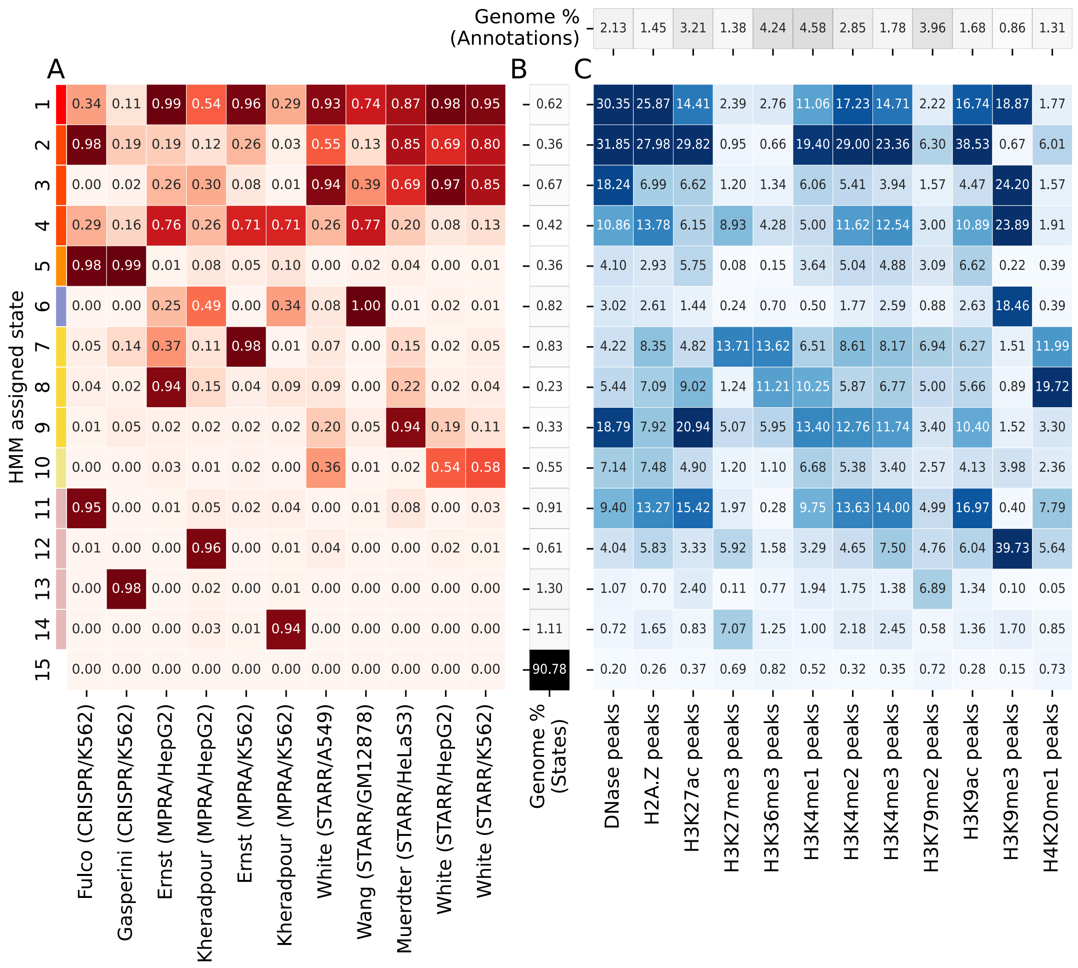


**Supplementary Figure S8: ChromScoreHMM emission parameters and chromatin mark peak enrichments for the 15-state model. (A)** Emission parameters of the ChromScoreHMM model (see Figure 3 for details). **(B)** Percentage of the genome assigned to each ChromScoreHMM state. **(C)** Fold enrichments for peak calls for individual chromatin marks based on imputed data from Roadmap Epigenomics. Top row indicates the percentage of the genome occupied by the peak call annotation. Enrichments and percentages are medians across cell types.

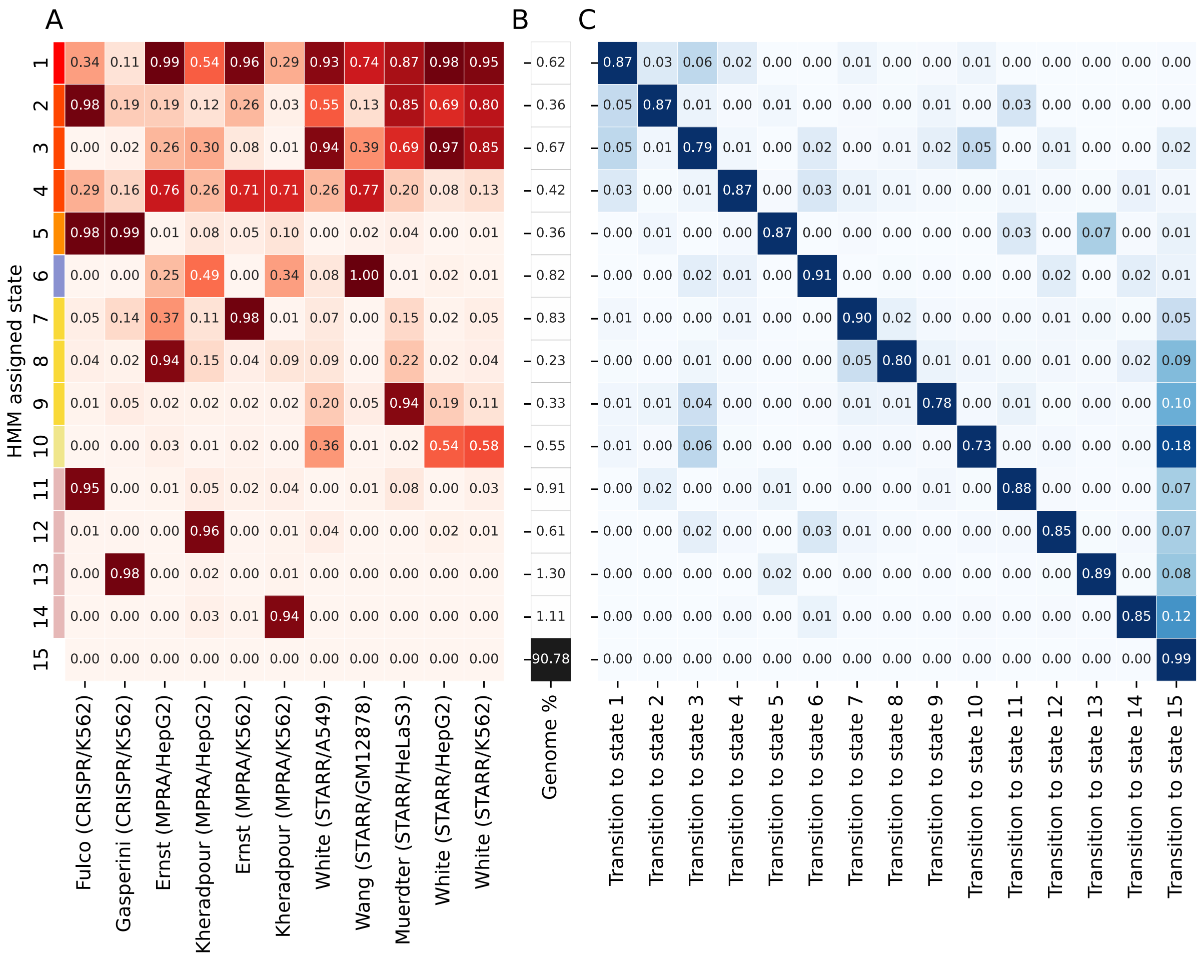


**Supplementary Figure S9: ChromScoreHMM emission and transition parameters for the 15-state model. (A)** Emission parameters of the model (see Figure 3 for details). **(B)** Percentage of the genome assigned to each ChromScoreHMM state. **(C)** Transition parameters of the model. Each value corresponds to the probability when in the state of the row of transitioning to the state of the column.

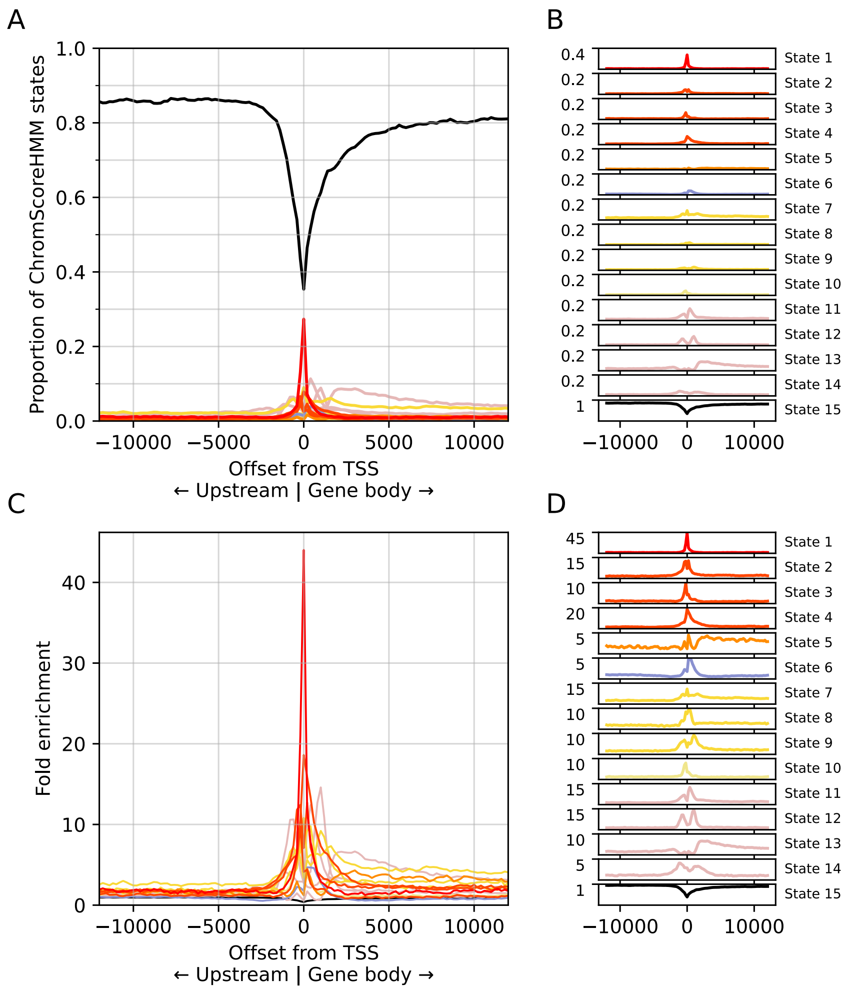


**Supplementary Figure S10: ChromScoreHMM state proportions around transcription start sites. (A)** Proportion of ChromScoreHMM state assignments represented by each ChromScoreHMM state as a function of genomic position centered around TSSs, averaged over cell types. See B for color legend. Negative offset values are upstream of the transcription start site and positive offset values are downstream. Offset zero refers to the TSS. **(B)** Individual plots of ChromScoreHMM state proportions and color legend. State 15 color changed from white to black for legibility. Bottom limits for the y-axes are 0 (omitted for clarity), upper limits as indicated on the plot. **(C)** Fold enrichments of ChromScoreHMM states as a function of genomic position, similar to (A). **(D)** Individual plots of ChromScoreHMM fold enrichments, similar to (B).

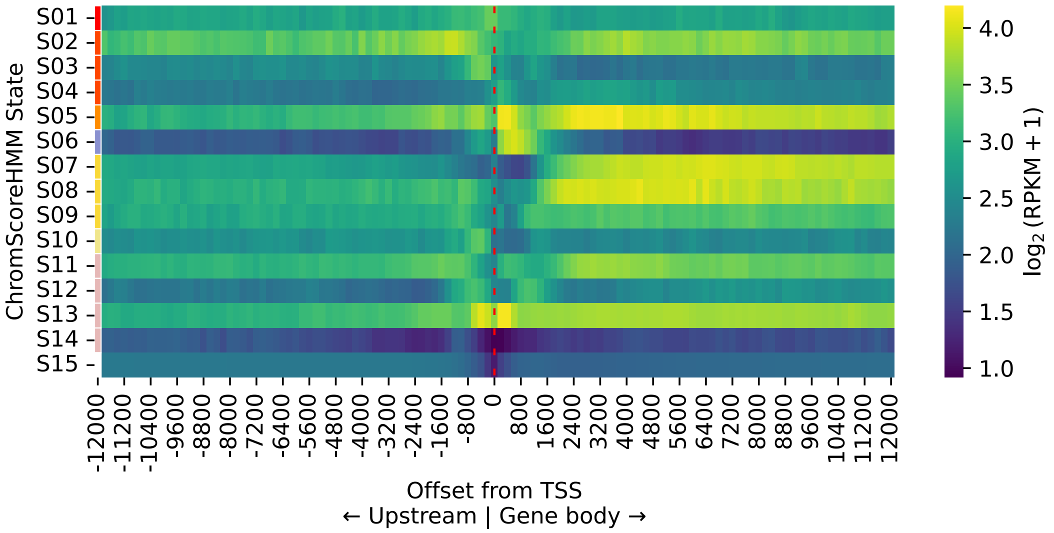


**Supplementary Figure S11: ChromScoreHMM gene expression patterns around transcription start sites.** Mean expression of nearby genes for positions within a +/-12 kb window around the TSSs (dashed red line) occupied by each ChromScoreHMM state. Each row corresponds to a ChromScoreHMM state and each column a position relative to a TSS. Expression values for each offset position and ChromScoreHMM state are the average $\log_2$(RPKM+1) values across both genes and cell types. RPKM: reads per kilobase of exon per million mapped reads.

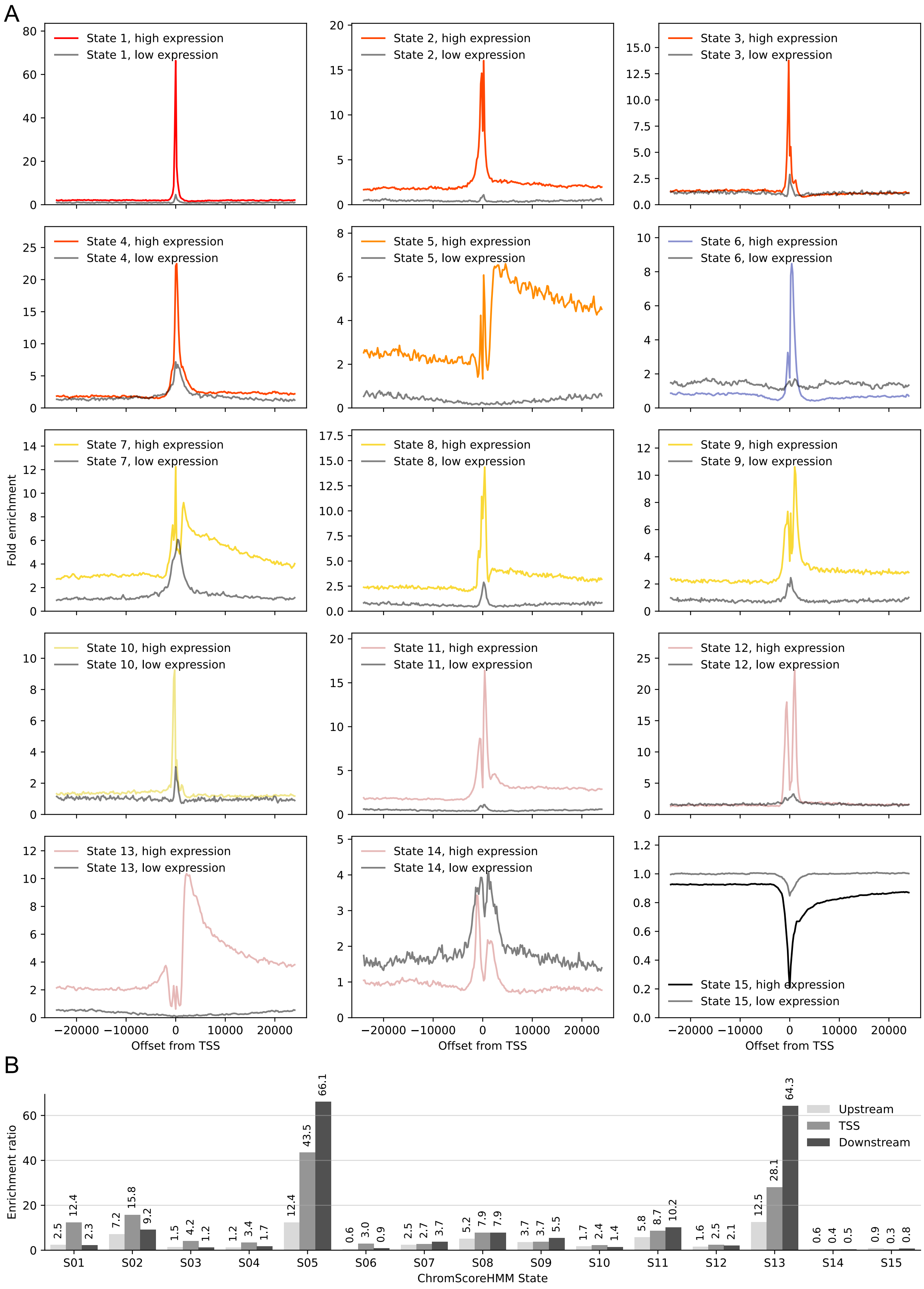
A
State 1, high expression
State 1, low expression
State 2, high expression
State 2, low expression
State 3, high expression
State 3, low expression
State 4, high expression
State 4, low expression
State 5, high expression
State 5, low expression
State 6, high expression
State 6, low expression
State 7, high expression
State 7, low expression
State 8, high expression
State 8, low expression
State 9, high expression
State 9, low expression
State 10, high expression
State 10, low expression
State 11, high expression
State 11, low expression
State 12, high expression
State 12, low expression
State 13, high expression
State 13, low expression
State 14, high expression
State 14, low expression
State 15, high expression
State 15, low expression
Fold enrichment
−20000 −10000 0 10000 20000
Offset from TSS
B
Enrichment ratio
Upstream
TSS
Downstream
S01 2.5 12.4 2.3
S02 7.2 15.8 9.2
S03 1.5 4.2 1.2
S04 1.2 3.4 1.7
S05 12.4 43.5 66.1
S06 0.6 3.0 0.9
S07 2.5 2.7 3.7
S08 5.2 7.9 7.9
S09 3.7 3.7 5.5
S10 1.7 2.4 1.4
S11 5.8 8.7 10.2
S12 1.6 2.5 2.1
S13 12.5 28.1 64.3
S14 0.6 0.4 0.5
S15 0.9 0.3 0.8
ChromScoreHMM State

**Supplementary Figure S12: ChromScoreHMM overlap enrichments around TSSs of high expression and low expression genes. (A)** Each subplot corresponds to a ChromScoreHMM state and shows the fold enrichments relative to the TSS for both a set of high expression and low expression genes as indicated by the color legend in each plot. High expression color corresponds to ChromScoreHMM state colors (Supplementary Figure S2). Negative offset values are upstream of the transcription start site and positive offset values are downstream. Offset zero refers to the TSS. Gene expression thresholds for high and low expression genes are $\log_2(\text{RPKM}+1) > 1$ and $\log_2(\text{RPKM}+1) < 0.01$, respectively. **(B)** Ratio of enrichments for high expression relative to low expression genes for each ChromScoreHMM state restricted to positions upstream of the TSS, at the TSS, or downstream of the TSS. The enrichment ratios correspond to the number of bases covered by a ChromScoreHMM state within an interval around high expression genes over the number of bases covered within the equivalent interval around low expression genes, each normalized by the total number of bases in the corresponding high expression and low expression gene intervals before computing the ratios. Upstream and downstream ratios computed over the 0.5-10 kb interval centered around the TSS. TSS ratio computed over the interval -200 to 200 bp.

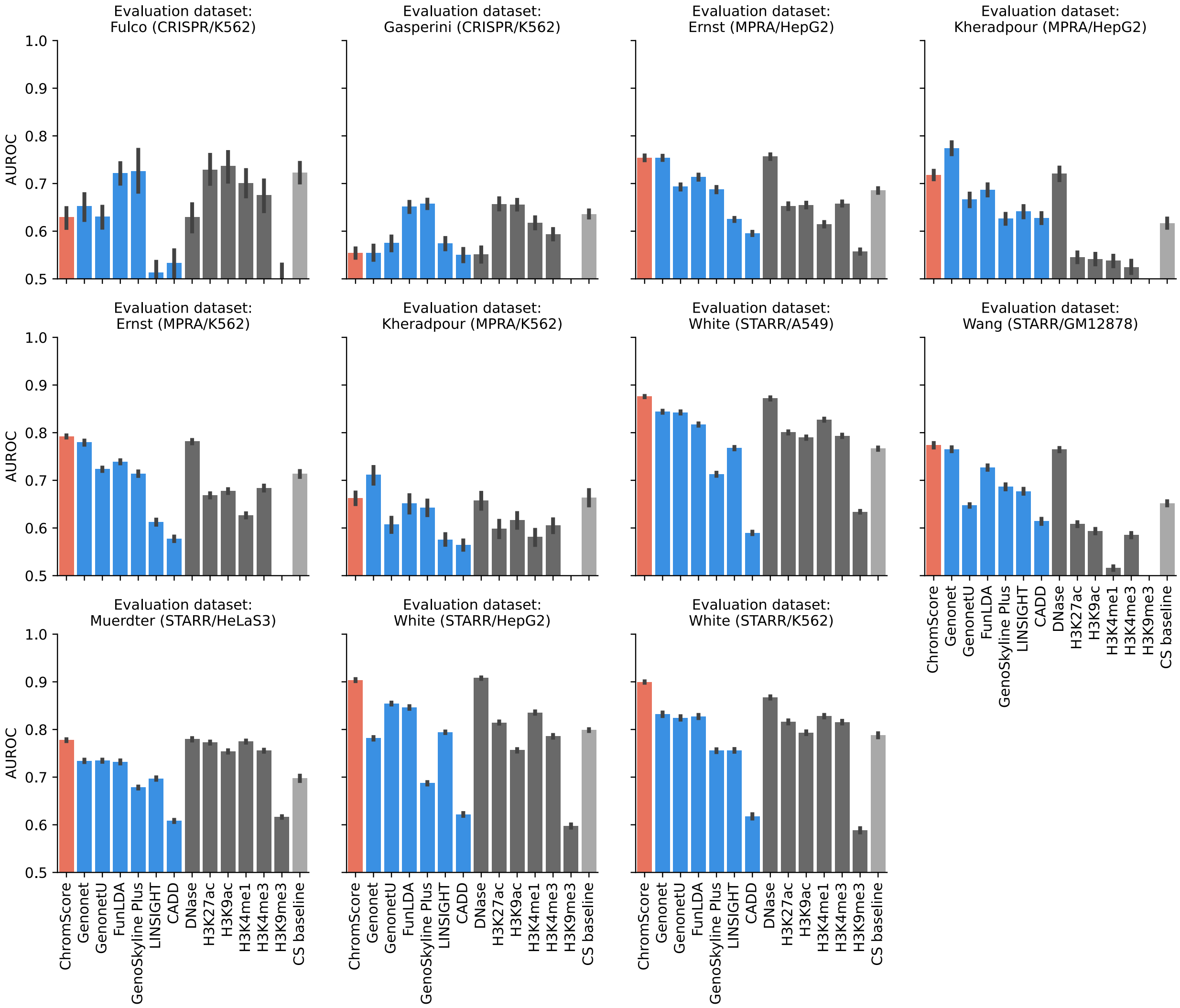


**Supplementary Figure S13: Predictive performance of ChromScore, external models and baselines.** Each panel corresponds to a functional characterization dataset used for evaluating regulatory activity predictions. ChromScore excludes all training data from evaluation cell types or from the same genomic position. Error bars indicate standard error across test partitions. AUROC: Area under receiver operator characteristic curve. Red bar: ChromScore, blue bars: external scores (Genonet, GenonetU, FunLDA, GenoSkylinePlus, LINSIGHT, CADD), dark gray bars: individual chromatin mark signals, light gray bar: chromatin state baseline model.

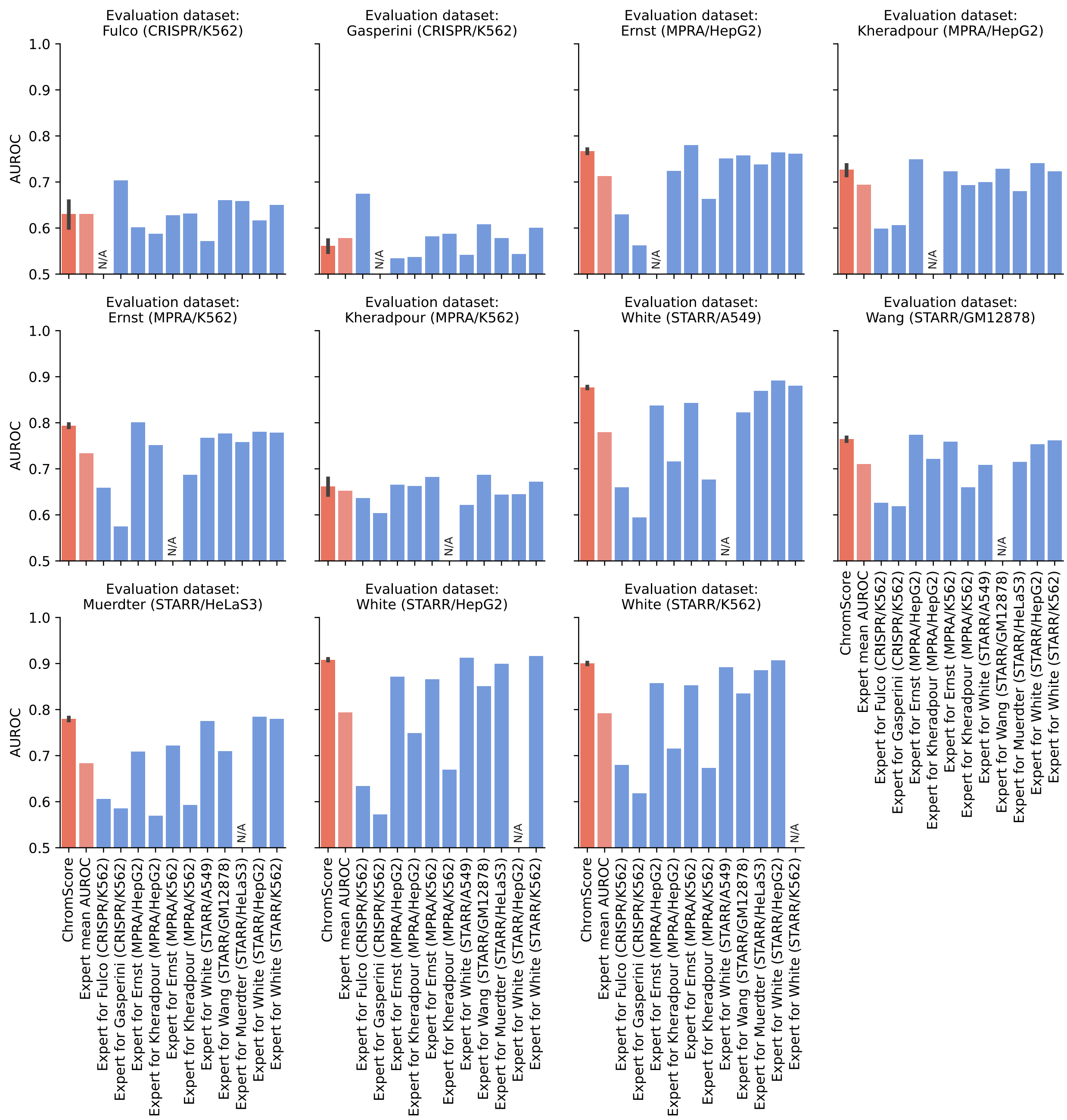


**Supplementary Figure S14: Predictive performance of ChromScore and individual experts.** Each panel corresponds to a functional characterization dataset used for evaluating regulatory activity predictions. Expert models trained on a given dataset were not evaluated on the same dataset and were marked “N/A”. See Supplementary Figure S4 for performance of expert models on the same dataset under cross-validation. In evaluating ChromScore, we excluded ChromScore all training data from evaluation cell types or from the same genomic position (Figure 4B, Methods). Error bar indicates standard error across ChromScore test partitions. AUROC: Area under receiver operator characteristic curve. Red bar: ChromScore, light red bar: mean AUROC of individual experts for the given evaluation, blue bars: individual experts.

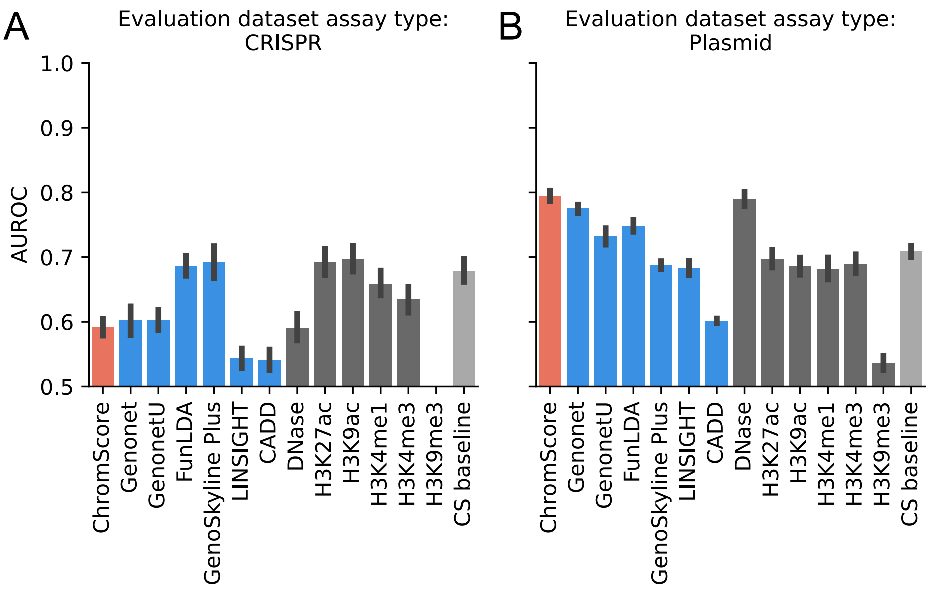


**Supplementary Figure S15: Predictive performance of ChromScore, external models and baselines in CRISPR and plasmid-based functional characterization datasets. (A)** Mean score performance evaluated in CRISPR-based datasets. **(B)** Mean score performance evaluated in plasmid-based datasets. Error bars indicate standard error across evaluations. AUROC: Area under receiver operator characteristic curve. Red bar: ChromScore, blue bars: external scores (Genonet, GenonetU, FunLDA, GenoSkylinePlus, LINSIGHT, CADD), dark gray bars: individual chromatin mark signals, light gray bar: chromatin state baseline model.

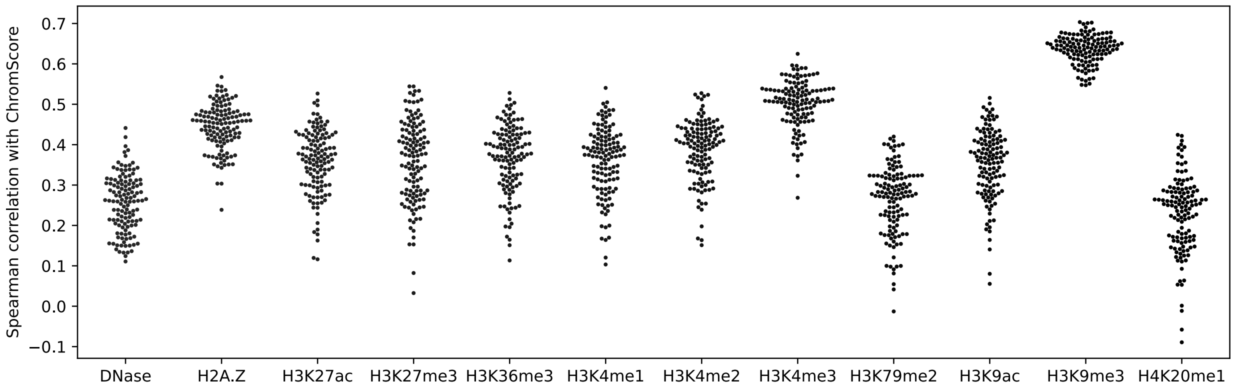


**Supplementary Figure S16: Spearman correlations between ChromScore and chromatin mark signal tracks.** Each point indicates correlation between the indicated mark and ChromScore in one cell type.

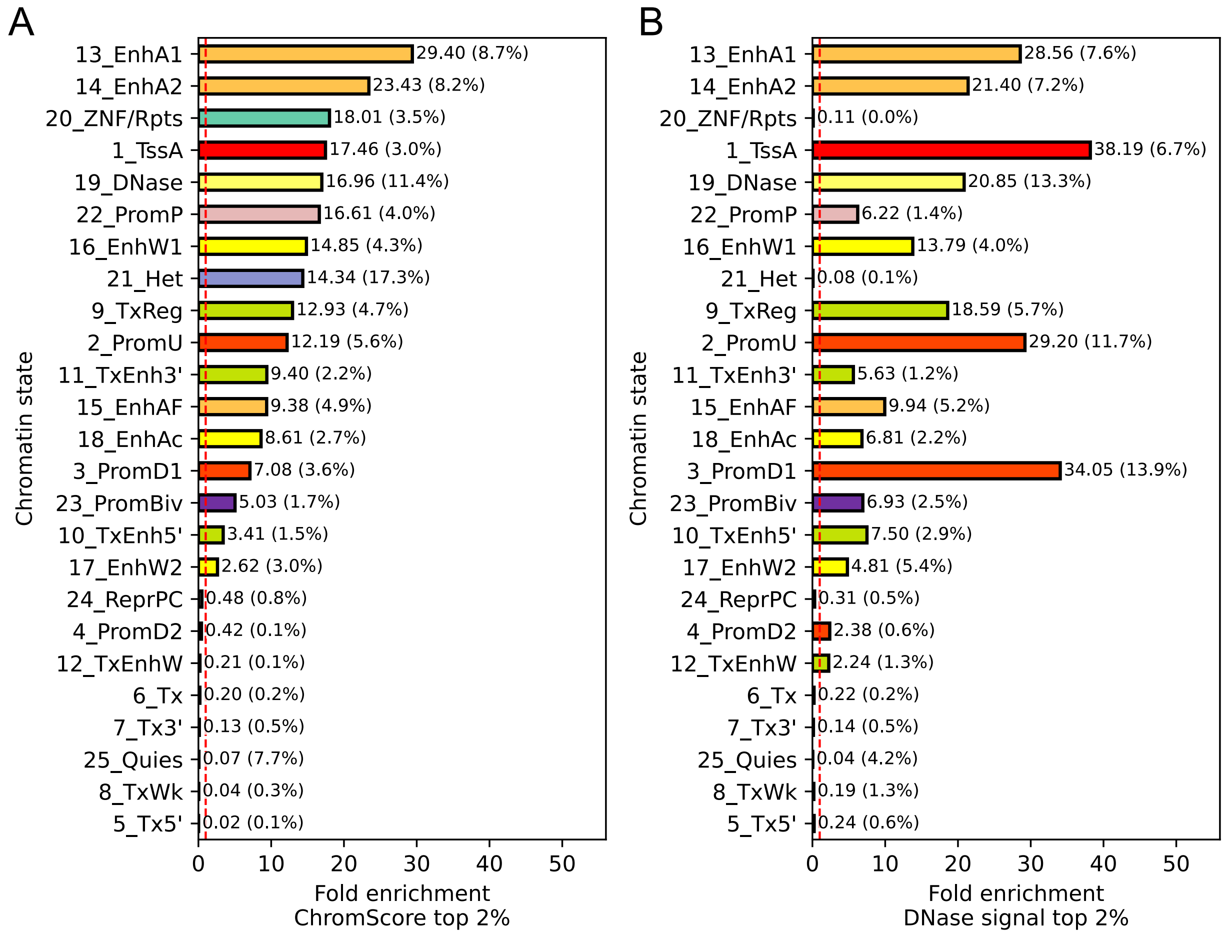


**Supplementary Figure S17: Chromatin state fold enrichments of top ChromScore and DNase-I hypersensitive regions.** Chromatin state fold enrichments for genomic regions assigned the top 2% of **(A)** ChromScore, **(B)** DNase-I hypersensitivity signal, geometric mean over cell types. Dashed red line indicates fold enrichment 1. Chromatin state composition average percentages within the top 2% of regions shown in parentheses. States are ordered as in **A** by ChromScore enrichment.

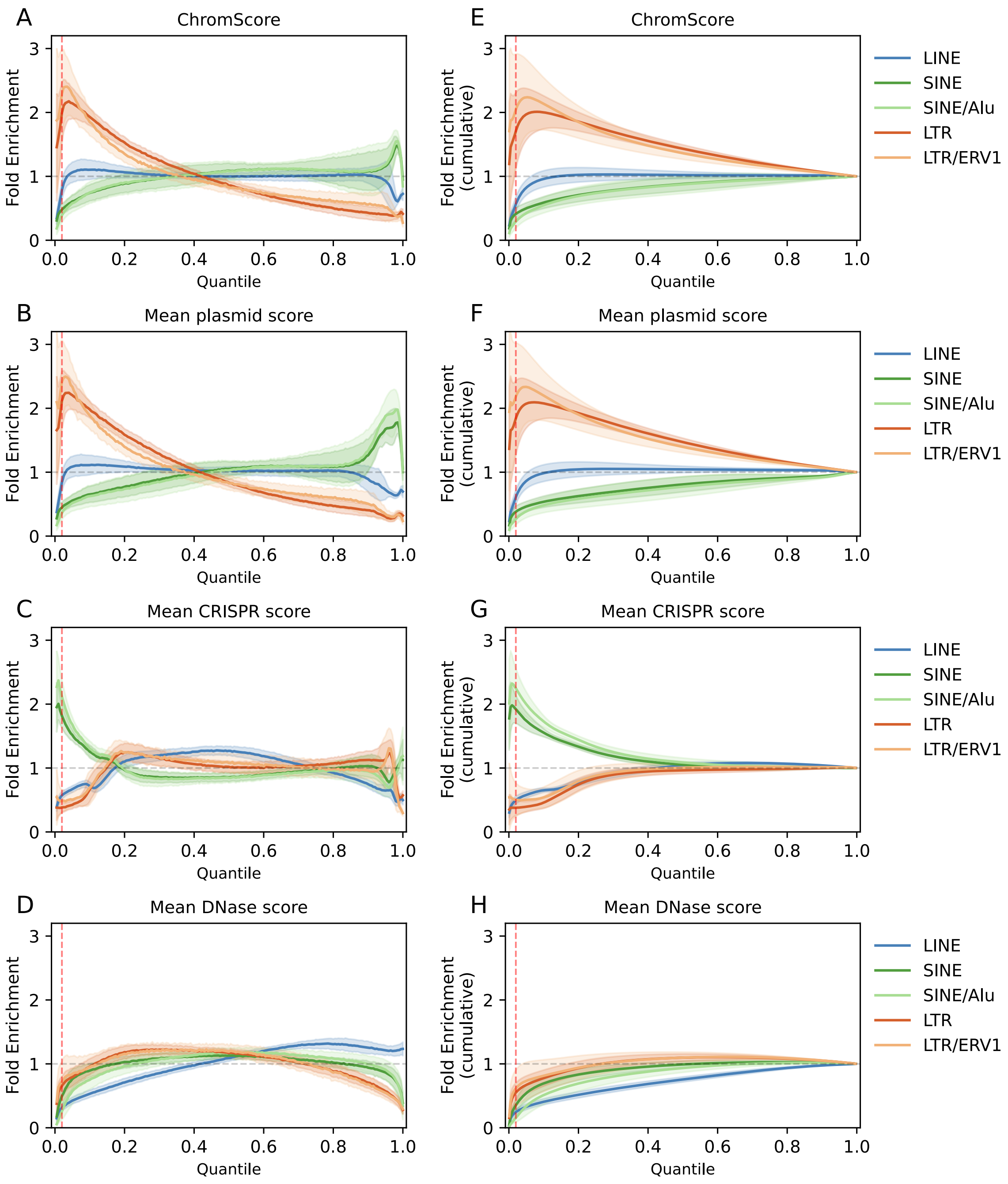


**Supplementary Figure S18: Repeat element fold enrichments for ChromScore, mean plasmid-based expert score, mean CRISPR-based expert score and DNase signal quantiles.** Fold enrichments for long interspersed nuclear elements (LINEs), all short interspersed nuclear elements (SINEs), the Alu subclass of SINEs, all long terminal repeats

(LTRs) and the endogenous retroviral sequence 1 (ERV1) subclass of LTRs, computed for **(A)** ChromScore, **(B)** mean plasmid-based expert score, **(C)** mean CRISPR-based expert score, **(D)** DNase signal, **(E-H)** same as (A-D) except for cumulative fold enrichments. Lines indicate mean enrichments across 127 cell types, shaded areas indicate standard deviations across cell types. Fold enrichments are computed relative to the genome background. Red vertical dashed line denotes the top 2% of scores. Gray horizontal line indicates fold enrichment of 1.

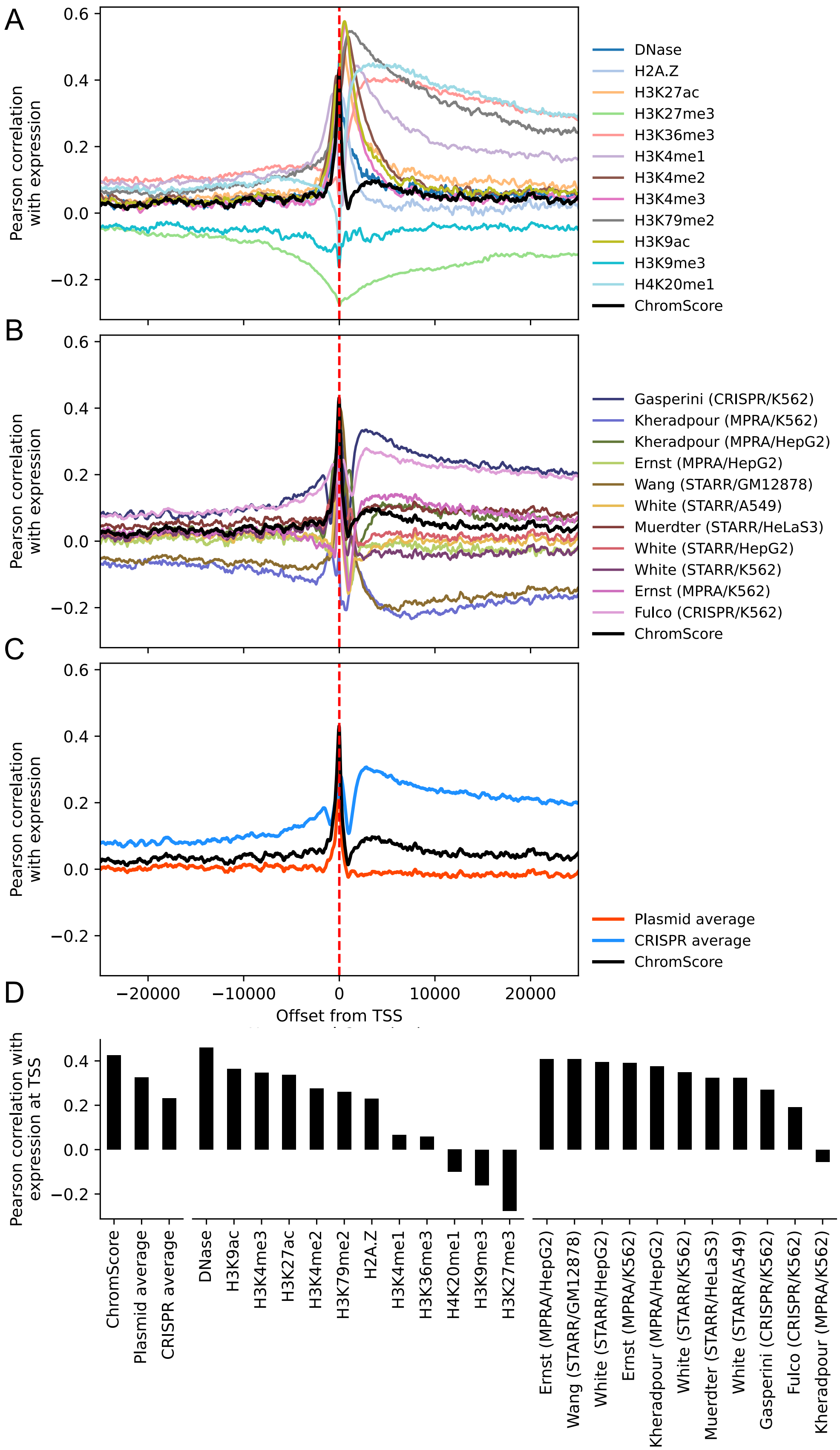
A
B
C
D
Pearson correlation with expression
Pearson correlation with expression at TSS
Offset from TSS
−20000
−10000
0
10000
20000
DNase
H2A.Z
H3K27ac
H3K27me3
H3K36me3
H3K4me1
H3K4me2
H3K4me3
H3K79me2
H3K9ac
H3K9me3
H4K20me1
ChromScore
Gasperini (CRISPR/K562)
Kheradpour (MPRA/K562)
Kheradpour (MPRA/HepG2)
Ernst (MPRA/HepG2)
Wang (STARR/GM12878)
White (STARR/A549)
Muerdter (STARR/HeLaS3)
White (STARR/HepG2)
White (STARR/K562)
Ernst (MPRA/K562)
Fulco (CRISPR/K562)
Plasmid average
CRISPR average
ChromScore
Plasmid average
CRISPR average
DNase
H3K9ac
H3K4me3
H3K27ac
H3K4me2
H3K79me2
H2A.Z
H3K4me1
H3K36me3
H4K20me1
H3K9me3
H3K27me3
Ernst (MPRA/HepG2)
Wang (STARR/GM12878)
White (STARR/HepG2)
Ernst (MPRA/K562)
Kheradpour (MPRA/HepG2)
White (STARR/K562)
Muerdter (STARR/HeLaS3)
White (STARR/A549)
Gasperini (CRISPR/K562)
Fulco (CRISPR/K562)
Kheradpour (MPRA/K562)

**Supplementary Figure S19: Cell type averages of Pearson correlations between gene expression and ChromScore, individual chromatin mark signals, expert scores.**

Correlation of gene expression ($log_2$(RPKM+1)) with ChromScore and (**A**) individual chromatin mark signals, **(B)** individual expert tracks, **(C)** and average of plasmid-based expert tracks and CRISPR-based expert tracks as a function of position relative to the TSS within a 24 kb window. **(D)** Bar graph reporting correlation with gene expression of A-C specifically at the TSS.

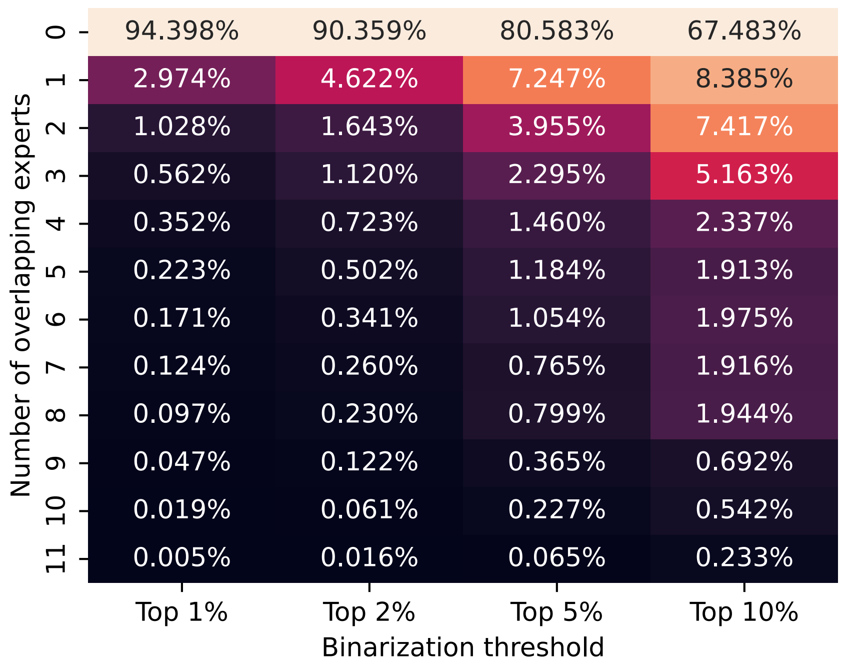


**Supplementary Figure S20: Percentage of genome above the binarization threshold by one or more experts.** The columns correspond to the binarization thresholds of top 1%, 2%, 5%, and 10% of the expert prediction scores. Each row corresponds to a number of experts. Values correspond to the percentage of 25 bp intervals in the genome for which exactly the number of experts of the row had scored it above the binarization threshold of the column.

**Supplementary Table 1: Overview of the functional characterization datasets.**

| Name | Cell type | Assay technology | Unit of observation | Citation | Number of activating loci | Number of neutral loci | Total number of data points |
|---|---|---|---|---|---|---|---|
| Fulco/K562 | K562 | CRISPR-dCas9 | Center nucleotide of target site | [5] | 69 | 747 | 816 |
| Gasperini/K562 | K562 | CRISPR-dCas9 | Center nucleotide of target site | [6] | 432 | 5122 | 5554 |
| Kheradpour/HepG2 | HepG2 | MPRA | Individual construct | [7] | 541 | 1548 | 2089 |
| Kheradpour/K562 | K562 | MPRA | Individual construct | [7] | 347 | 1742 | 2089 |
| Ernst/HepG2 | HepG2 | MPRA | Nucleotide with absolute maximum score in tiled region | [8] | 2405 | 10,894 | 13,299 |
| Ernst/K562 | K562 | MPRA | Nucleotide with absolute maximum score in tiled region | [8] | 2519 | 10,162 | 12,681 |
| Wang/GM12878 | GM12878 | STARR-seq | Center of driver elements | [9] | 2409 | 7227 | 9636 |
| Muerdter/HeLaS3 | HeLa-S3 | STARR-seq | Center of peak call | [10] | 9613 | 28,839 | 38,452 |
| White/A549 | A549 | STARR-seq | Center of peak call | [11] | 6929 | 20,787 | 27,716 |
| White/HepG2 | HepG2 | STARR-seq | Center of peak call | [12] | 5199 | 15,597 | 20,796 |
| White/K562 | K562 | STARR-seq | Center of peak call | [12] | 3571 | 10,713 | 14,284 |

**References**


1. Ernst J, Kellis M. ChromHMM: automating chromatin-state discovery and characterization. Nat Methods. 2012;9:215–6.

2. Ernst J, Kellis M. Large-scale imputation of epigenomic datasets for systematic annotation of diverse human tissues. Nat Biotechnol. 2015;33:364–76.

3. Roadmap Epigenomics Consortium, Kundaje A, Meuleman W, Ernst J, Bilenky M, Yen A, et al. Integrative analysis of 111 reference human epigenomes. Nature. 2015;518:317–30.

4. Ernst J, Kellis M. Chromatin-state discovery and genome annotation with ChromHMM. Nat Protoc. 2017;12:2478–92.

5. Fulco CP, Nasser J, Jones TR, Munson G, Bergman DT, Subramanian V, et al. Activity-by-contact model of enhancer–promoter regulation from thousands of CRISPR perturbations. Nat Genet. 2019;51:1664–9.

6. Gasperini M, Hill AJ, McFaline-Figueroa JL, Martin B, Kim S, Zhang MD, et al. A Genome-wide Framework for Mapping Gene Regulation via Cellular Genetic Screens. Cell. 2019;176:377-390.e19.

7. Kheradpour P, Ernst J, Melnikov A, Rogov P, Wang L, Zhang X, et al. Systematic dissection of regulatory motifs in 2000 predicted human enhancers using a massively parallel reporter assay. Genome Res. 2013;23:800–11.

8. Ernst J, Melnikov A, Zhang X, Wang L, Rogov P, Mikkelsen TS, et al. Genome-scale high-resolution mapping of activating and repressive nucleotides in regulatory regions. Nat Biotechnol. 2016;34:1180–90.

9. Wang X, He L, Goggin SM, Saadat A, Wang L, Sinnott-Armstrong N, et al. High-resolution genome-wide functional dissection of transcriptional regulatory regions and nucleotides in human. Nat Commun. 2018;9:5380.

10. Muerdter F, Boryń ŁM, Woodfin AR, Neumayr C, Rath M, Zabidi MA, et al. Resolving systematic errors in widely used enhancer activity assays in human cells. Nat Methods. 2018;15:141–9.

11. White K. ENCSR895FDL. 2020.

12. Lee D, Shi M, Moran J, Wall M, Zhang J, Liu J, et al. STARRPeaker: uniform processing and accurate identification of STARR-seq active regions. Genome Biol. 2020;21:298.
